# The molecular mechanism of lipid uptake by membrane-anchored bridge-like lipid transfer proteins

**DOI:** 10.1101/2025.08.01.668188

**Authors:** Daniel Álvarez, Paige Chandran Blair, Cristian Rocha-Roa, Michael Davey, Elizabeth Conibear, Stefano Vanni

## Abstract

Lipid transport by bridge-like lipid transfer proteins (BLTPs) is emerging as a key process in lipid and cellular metabolism in both physiological and pathological conditions. However, the precise mechanism of lipid transport by BLTPs has remained elusive. Here, we use extensive all-atom molecular dynamics simulations to characterize the precise mechanism of lipid transfer into the BLTP hydrophobic cavity from donor membranes. For multiple BLTPs, we observe the ability to extract and solubilize lipids without lipid selectivity, and we identify membrane destabilization as a critical parameter to achieve effective lipid desorption. We rationally design a mutant BLTP with altered ability to destabilize lipid bilayers, and we show that this abolishes lipid desorption *in silico* and protein function *in vivo*. Taken together, our data provide an atomic-level description of the mechanism of lipid transport by BLTPs, ultimately suggesting alternative strategies to interfere with their activity.

## Introduction

Lipid membranes constitute the physical barriers that separate the cell from the external environment, and they also allow for intracellular compartmentalization by subdividing the cellular space into distinct organelles.^1,2^ As a consequence, they are essential for cell survival not only due to their protective role, but also because of their direct involvement in many cellular trafficking processes. To maintain such a complex infrastructure and to guarantee correct cellular homeostasis, lipids need to be constantly transported between organelles in a fast and efficient manner.^3^

Recent compelling evidence indicates that most lipids are transported by non-vesicular lipid transfer processes that mainly occur at membrane contact sites (MCSs) via lipid transfer proteins (LTPs).^4–6^ There are hundreds of known LTPs in all organisms, but most of them are relatively small, with a cavity that can accommodate only one or two lipids. For those proteins, the proposed lipid transport mechanism is the so-called shuttle mechanism, which consists of the solubilization of a lipid inside the LTP cavity from a donor membrane, followed by its delivery to an acceptor membrane.^5–7^ However, this form of transport is not sufficient to explain fast bulk lipid transport under stress conditions.

During the past few years, a new family of LTPs has been discovered, the bridge-like LTPs (BLTPs).^6,8^ These proteins share a common shape, consisting of a long (∼10-30 nm) hydrophobic tunnel, surrounded by hydrophilic residues that face the protein exterior. Consequently, a new type of lipid transport mechanism has been proposed, in which BLTPs connect two membranes within a MCS so that lipids can directly flow from one membrane to another through the protein hydrophobic cavity.^6,9^ Since the first discovery of the lipid transfer activity of BLTPs, these proteins have been identified in a wide variety of organisms, targeting different organelles and playing an important role in lipid homeostasis and membrane biogenesis.^8,10,11^ Their importance is highlighted by their presence at almost all known MCSs, and their extensive involvement in neurological disorders, including Parkinson’s disease, Cohen syndrome, ataxia, or chorea-acanthocytosis, among others.^12–18^

Due to their large size (∼150-550 kDa) and extensive hydrophobic character, the reconstitution and determination of the experimental structure of these proteins has posed a major challenge. Despite these difficulties, structural insights into the structure and function of BLTPs, albeit sparse, are available: several cryo-EM low-resolution structures of BLTPs have demonstrated the general tube shape of this protein family, with long cylindrical hydrophobic cavities, and their relative orientation to membranes,^19–21^ whereas high resolution structures of BLTP fragments from X-ray diffraction and cryo-EM have unveiled the features of the actual domains that form the hydrophobic cavity as well as the orientation of the lipids inside them,^22–24^ including in a recent high-resolution cryo-EM structure of a long N-terminal fragment of the ER-anchored BLTP LPD-3 bound to multiple lipids.^24^

These studies, together with the analysis of full-length AI-predicted structures of BLTPs, have confirmed that these proteins consist of a long hydrophobic groove formed by repeating beta groove (RBG) domains and a hydrophilic exterior that maintains the solubility within the cytosol. ^25,26^ They also typically possess amphipathic helices at their extremes, which may play a role in membrane targeting,^27–29^ while a few of them possess a transmembrane (TM) domain at the N-terminal part of the protein.

Although the structural knowledge of BLTPs has largely improved in recent years, the details of their lipid transport mechanism remain a mystery. *In vitro* experiments have demonstrated the ability of BLTPs to solubilize and transport lipids^22,23,30–32^, and mutation strategies in which hydrophobic residues inside the cavity are exchanged for charged residues have proved successful in abolishing lipid transport and protein function^32–36^. However, the proteins in these experiments are often not directly attached to both bilayers, so a shuttle mechanism can still not be totally discarded. Moreover, even lipid-bound BLTP structures only provide static information about the lipid transport mechanism, and, due to the use of detergents, only in the absence of native-like membrane conditions. Therefore, many questions regarding BLTPs remain unanswered, including the lipid uptake mechanism, lipid flow and the driving forces of the process.^37^

In this work, we use all-atom (AA) molecular dynamics (MD) simulations to describe with molecular detail the lipid uptake mechanism of several BLTP fragments containing TM domains. Our simulations demonstrate the ability of all studied BLTPs to extract and “solubilize” lipids without lipid selectivity and provide valuable insights into the mechanisms and energetics of lipid desorption. Based on the molecular understanding derived from the simulations, we design a new strategy to abolish their function by mutating residues located at the protein-membrane interface rather than inside the hydrophobic tunnel. Taken together, our data provide an atomic-level description of the mechanism of lipid transport by BLTPs.

## Results

### BLTPs can spontaneously desorb multiple lipids from lipid bilayers

To investigate the mechanism of lipid transport by BLTPs at the molecular level, we opted to use AA MD simulations, as this technique is well-suited to investigate dynamic protein-membrane and protein-lipid interactions at the molecular scale.^38–42^ As a model system, we chose to initially work with the yeast BLTP Csf1 for two main reasons. First, at its N-terminus, this protein possesses a TM domain that anchors it to the ER membrane.^43^ This structural property is shared by other BLTPs (Csf1 homologues such as LPD-3 or BLTP1, and Hobbit/BLTP2 proteins) but it is not a general feature of this protein family, as other members, including Vps13/VPS13 and Atg2/ATG2, lack this domain. However, from a modelling point of view, the presence of the TM domain greatly facilitates the correct positioning of the protein within the ER membrane (Fig. 1a).

**Figure 1.**
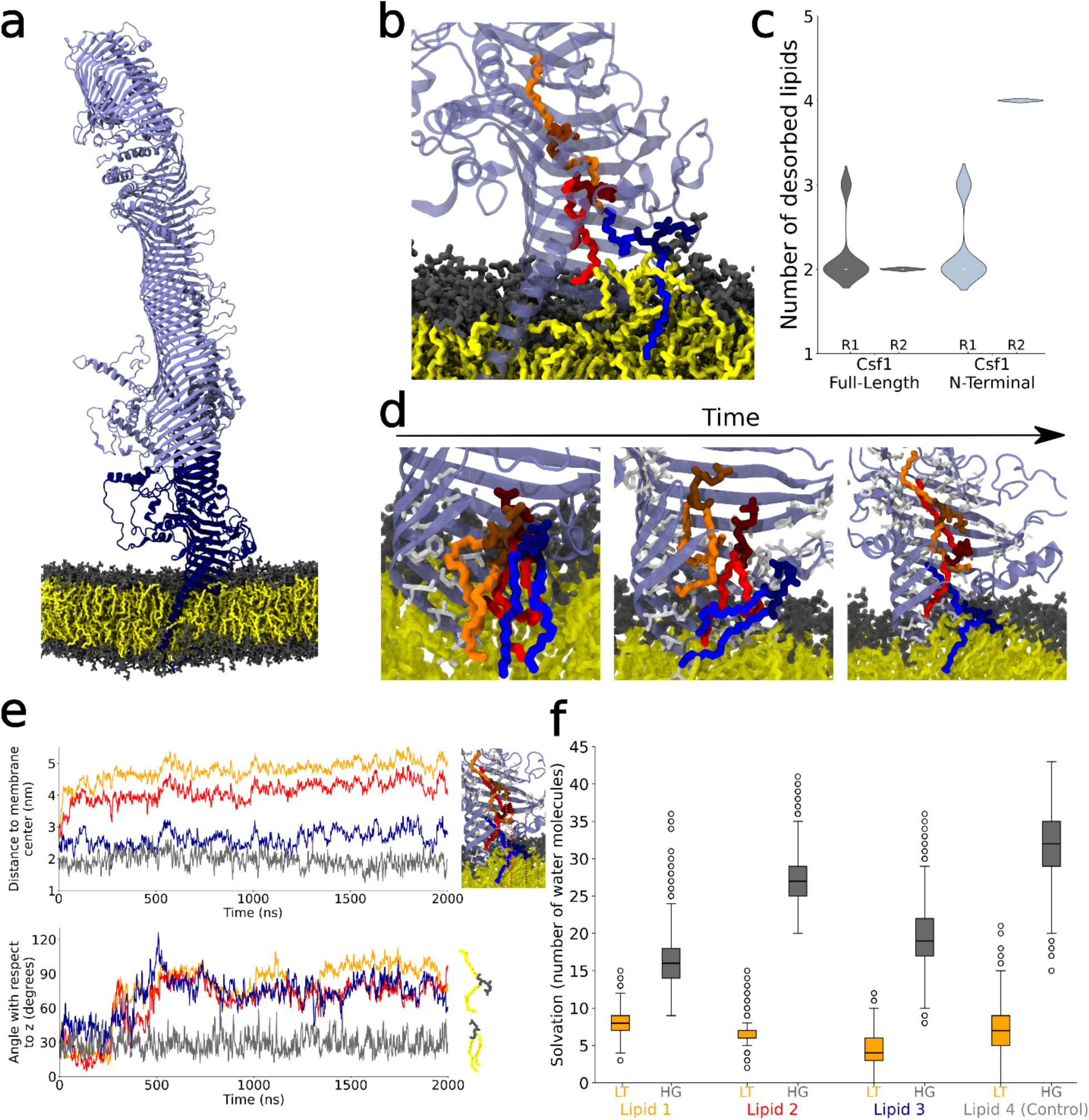
BLTP can spontaneously desorb multiple lipids from lipid bilayers. **a.** Full-length model of Csf1 embedded in a DOPC lipid bilayer, with the N-terminal segment of Csf1 considered in our subsequent AA MD simulations highlighted in darker blue (residues 1-645). **b.** Snapshot of the final frame of the full-length AA MD simulation of Csf1, showing the desorbed lipids within the protein cavity. **c** Violin plots of the number of desorbed lipids in the two replicas (R1, R2) of the full-length and the N-terminal AA MD simulations of Csf1. **d**. Representative snapshots of the mechanism of lipid uptake by Csf1. Desorbed lipids are coloured in orange, red, and blue, hydrophobic residues in contact with the tails of the desorbed lipids are depicted in white. **e.** Top: Time traces of the distance along *z* from the membrane center of the P atoms of the lipids depicted in panel b, with the same colour scheme, as well as of a control lipid that stays in the membrane at all times (grey colour lines); Bottom: Time traces of the average angle of their lipid tails with respect to the membrane normal. **f.** Boxplots of solvation (number of water molecules at a close distance) for the lipid tails (LT, orange boxplots) and headgroups (HG, grey boxplots) of the three lipids depicted in panel b as well as of a control lipid.

Second, among the BLTPs containing TM domains, Csf1 has been the most extensively studied, with several published works that have unveiled its role in the biosynthesis of GPI anchors and in the homeoviscous adaption of yeast under cold conditions.^43,44^ In addition, the AlphaFold2 (AF) predicted structures of Csf1 and its orthologs have high pLDDT scores, and the AF model of its *C. elegans* ortholog, LPD-3, showed very good agreement with the structure solved using single particle cryo-EM.^24,45^ The human homologue of Csf1, BLTP1, is also of great interest, as it has been linked to a rare neurological disease known as Alkuraya-Kucinskas syndrome.^46^

As a first step, we focused on the mechanism of lipid uptake by Csf1 in a simple membrane model. To do so, we performed AA MD simulations of Csf1 embedded in a pure DOPC lipid bilayer (Fig. 1a) through its TM domain. After a short simulation time, we observed the spontaneous desorption of multiple lipids in both replicas (Fig. 1b, c), with 2 lipids entering the hydrophobic cavity of Csf1 in each case (Fig. 1b, c, Supp. Movie S1).

Since lipid desorption is exclusively localized in the membrane-proximal N-terminal domain of Csf1 (Fig. 1b, Fig. S1), and given the large size of the protein, we next decided to use a smaller model system, and we switched to MD simulations of the N-terminal part of Csf1 (residues 1 to 645, Fig. 1a). Also in the smaller model system, we observed spontaneous desorption of multiple lipids in unbiased AA MD simulations, comparable to that of the full-length protein (Fig. 1c, d; Supp. Movie S2). Once desorbed, the lipids diffuse quite freely inside the protein cavity, moving to a distance of up to 5 nm from the membrane center (Fig. 1e). In addition to the desorbed lipids, a few lipid tails insert into the protein cavity, interacting with both the desorbed lipids and the protein hydrophobic residues (Fig. 1d, Fig. S1).

Mechanistically, in our AA simulations we observe that the desorbed lipids first align themselves with the hydrophobic cavity, keeping the lipid tails close to the hydrophobic residues of the beta-sheets of the first RBG domain, which is partially inserted into the membrane, while the headgroups remain solvated thanks to the opening of the groove. Then, the headgroup of the first desorbed lipid (orange lipid in Fig. 1d) displaces upwards, away from the membrane centre, and the lipid tails start interacting with a wider range of protein hydrophobic residues. Consequently, the second lipid (red lipid in Fig. 1d), is also drawn up to maintain its interactions with the first lipid. In both cases, the lipids keep the headgroups facing away from the beta sheets. A third lipid (blue lipid in Fig. 1d) at the entry of the cavity adopts a more perpendicular orientation while still in the membrane, with one of its lipid tails pointing upwards to interact with the first two desorbed lipids. Finally, the orientation of the upper lipid changes, adopting an open conformation with the lipid tails facing opposite directions, while keeping fixed the position of the headgroup (Fig. 1d, e, Fig. S1). Again, this triggers the same conformational change in the second lipid, which also extends one of its lipid tails towards the first lipid. Due to the displacement of the lipid tails of these lipids, lipid displacement at the entry of the cavity becomes more pronounced as well. Consequently, the average angle of the lipid tails with respect to the membrane normal is almost perpendicular for the three lipids (Fig. 1e, Fig. S2), while concomitantly allowing the solvation of the highly hydrophilic headgroups (Fig. 1f). Once these events occur, the lipids reach a metastable state. In this state, lipid tail solvation for the desorbed lipids remains similar to that of bulk membrane lipids thanks to the simultaneous interaction with both the hydrophobic residues of the protein and with the other lipids in the cavity (Fig. 1f).

### Lipid uptake by BLTPs is not lipid specific

Next, we investigated whether Csf1 displays selectivity for certain lipids in our desorption simulations. To do so, and to also discard possible artifacts from using a pure DOPC membrane, we performed AA simulations with different membrane compositions: 1. a pure DOPE membrane, 2. a membrane with heterogeneous composition and highly negatively charged lipids that contains DOPC, DOPE, PI3P, DOPS and ergosterol in a 40:25:25:10:10 ratio, and 3. an ER-like membrane with the same composition ratio as in ^47^, that is 39.4 % PC, 12.1 % PE, 41.7 % PI, 3.8 % PS, 3.0 % PA of phospholipids, and 4.7 % of ergosterol (see Table S1).

In all cases, we again observed that several lipids desorb from the membrane to the hydrophobic cavity of the protein (Fig. 2a, b), with no apparent lipid specificity (Table 1) and with a mechanism that is similar to the one described for the DOPC membrane (Fig. S1, Fig. S2). It is noteworthy that even PI3P lipids, with a high negative charge and large polar headgroup, can be taken up by Csf1 in a relatively short simulation time, as the headgroups always face the opened part of the protein and thus remain solvated, away from the hydrophobic residues present in the beta-sheets of Csf1 (Fig. S2). Although there is not any observed specificity regarding the uptake of phospholipids, we could not observe the desorption of ergosterol during our MD simulations. However, the concentration of this lipid is much lower compared to that of the other lipids in our simulations, so we cannot completely rule out that BLTPs could also desorb sterols within their cavity.

**Figure 2.**
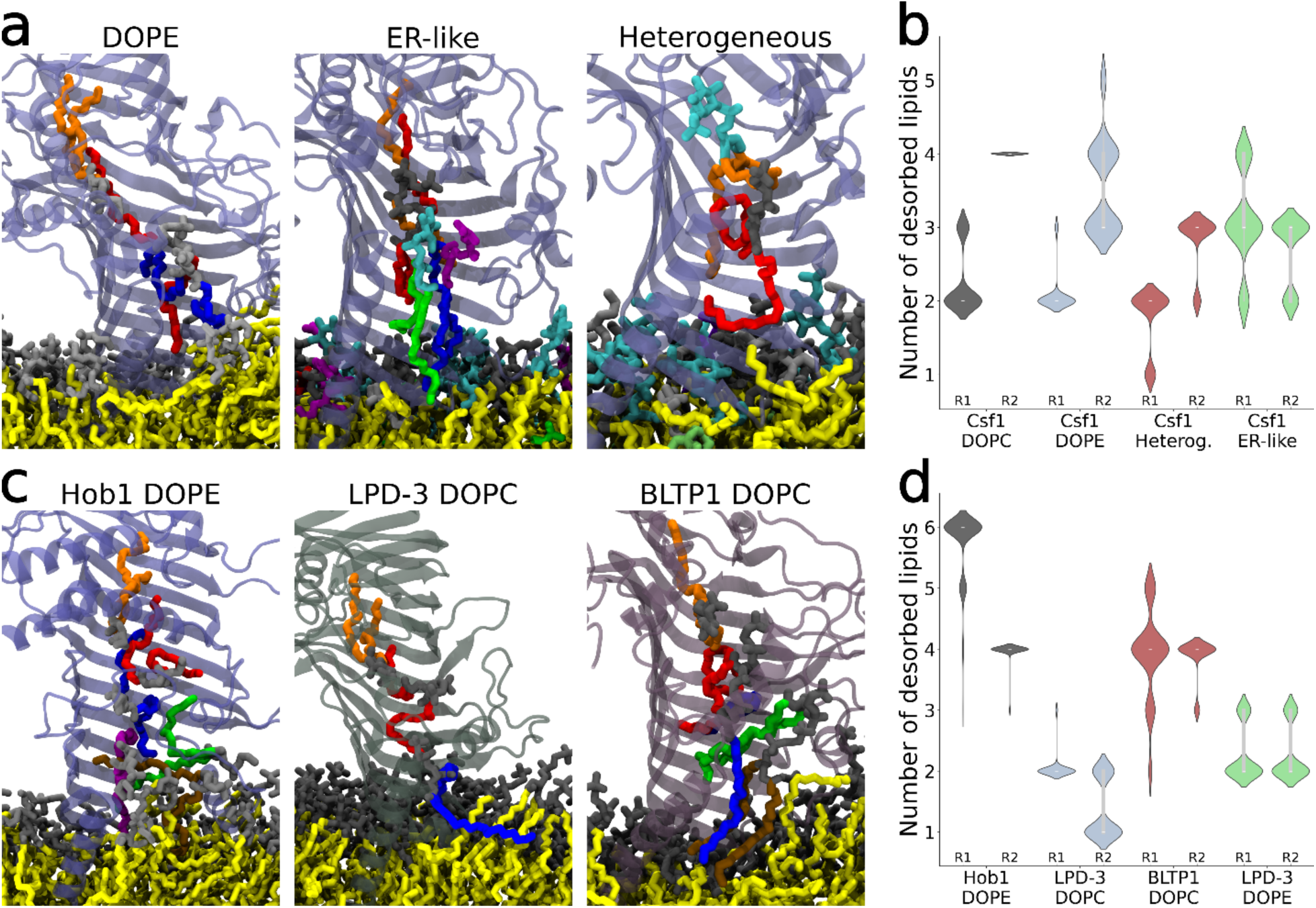
BLTPs uptake lipids with no specificity. **a.** Renders of the final frames of the AA MD simulations of Csf1 embedded in a DOPE, an ER-like membrane, and a membrane with heterogeneous composition and highly negative lipids. The headgroups of PE lipids are depicted in gray, of PC lipids in black, of PS in violet, of PA lipids in blue and of PI lipids in brown. Ergosterol is coloured in green. **b.** Violin plots of the number of desorbed lipids with each of the membrane compositions (R1 and R2 refer to replicas 1 and 2, respectively) **c.** Renders of the final frames of the AA MD simulations of Hob1, LPD-3 and BLTP1 showing the desorption of multiple lipids **d.** Violin plots of the number of desorbed lipids with each of the tested BLTPs (R1 and R2 refer to replicas 1 and 2, respectively)

**Table 1.** List of simulated BLTPs embedded in lipid bilayers along with the simulation time and the nature of the desorbed lipids.

| Protein | Composition | Simulation time | Desorbed lipids |
| --- | --- | --- | --- |
| Csf1 (full-length) | DOPC | 1 μs, 1 μs | DOPC |
| Csf1 (1-645) | DOPC | 2 μs, 1 μs | DOPC |
| Csf1 (1-645) | DOPE | 1 μs, 1 μs | DOPE |
| Csf1 (1-645) | Heterogeneous | 1 μs, 1 μs | PI3P, DOPC |
| Csf1 (1-645) | ER-like | 2 μs, 1 μs | DYPC, YOPC, PYPI, POPS |
| Csf1 F80R L99E (1-645) | DOPC | 1 μs, 1 μs | - |
| LPD-3 (1-578) | DOPC | 2 μs, 1 μs | DOPC |
| LPD-3 (1-578) | DOPE | 1 μs, 1 μs | DOPE |
| BLTP1 (1-594) | DOPC | 2 μs, 1 μs | DOPC |
| Hob1 (1-484) | DOPE | 2 μs, 1 μs | DOPE |

### Multiple BLTPs containing a transmembrane domain uptake lipids spontaneously

Finally, to exclude that what we observed is a Csf1-specific behaviour, we tested whether other BLTPs with a TM domain at their N-terminus would also spontaneously desorb lipids from the membrane. We thus selected Csf1 orthologs in other species: LPD-3 (in *C. elegans*) and BLTP1 (in humans), as well as a BLTP2-like protein (Hob1, also known as Fmp27, in *S. cerevisiae*), and embedded their TM-helix N-terminal fragment in either DOPC or DOPE membranes (Table 1). In all tested simulations we observed the uptake of several lipids into the BLTPs’ cavity (Fig. 2c, d and Fig. S1), with a desorption mechanism again similar to the one of Csf1 (Fig. S1, Fig. S2). Therefore, we can conclude that the observed phenomena of spontaneous lipid uptake by Csf1 can be extrapolated to all BLTPs with TM domains and to membranes with heterogeneous composition.

### Lipid uptake is driven by the replacement of within-bilayer lipid-lipid with lipid-protein hydrophobic contacts

The desorption of lipids from a membrane is an unfavourable process due to the breaking of multiple lipid-lipid hydrophobic interactions.^48^ Hence, the observed desorption of lipids from the membrane to the hydrophobic cavity of Csf1 in our AA simulations points to a potential compensation of this energetic loss via the establishment of energetically favourable interactions of the desorbed lipids with the protein.

To quantify the extent to which the protein affects membrane properties to promote lipid uptake, we first computed the local membrane curvature in the protein proximity (Fig. 3a, Fig. S4). By protruding into the membrane bilayer at the entry of the cavity, Csf1 generates positive curvature at this region (Fig. 3a, Fig. S3). Additionally, calculation of hydrophobic lipid-lipid contacts at the membrane (Fig. 3b) shows that the protein decreases lipid packing in its proximity. In agreement with this, the angle of the lipid tails with respect to the membrane normal is more perpendicular in the vicinity of this region (Fig. 3c), which is indicative of the formation of protein-lipid hydrophobic interactions at the membrane interface.

**Figure 3.**
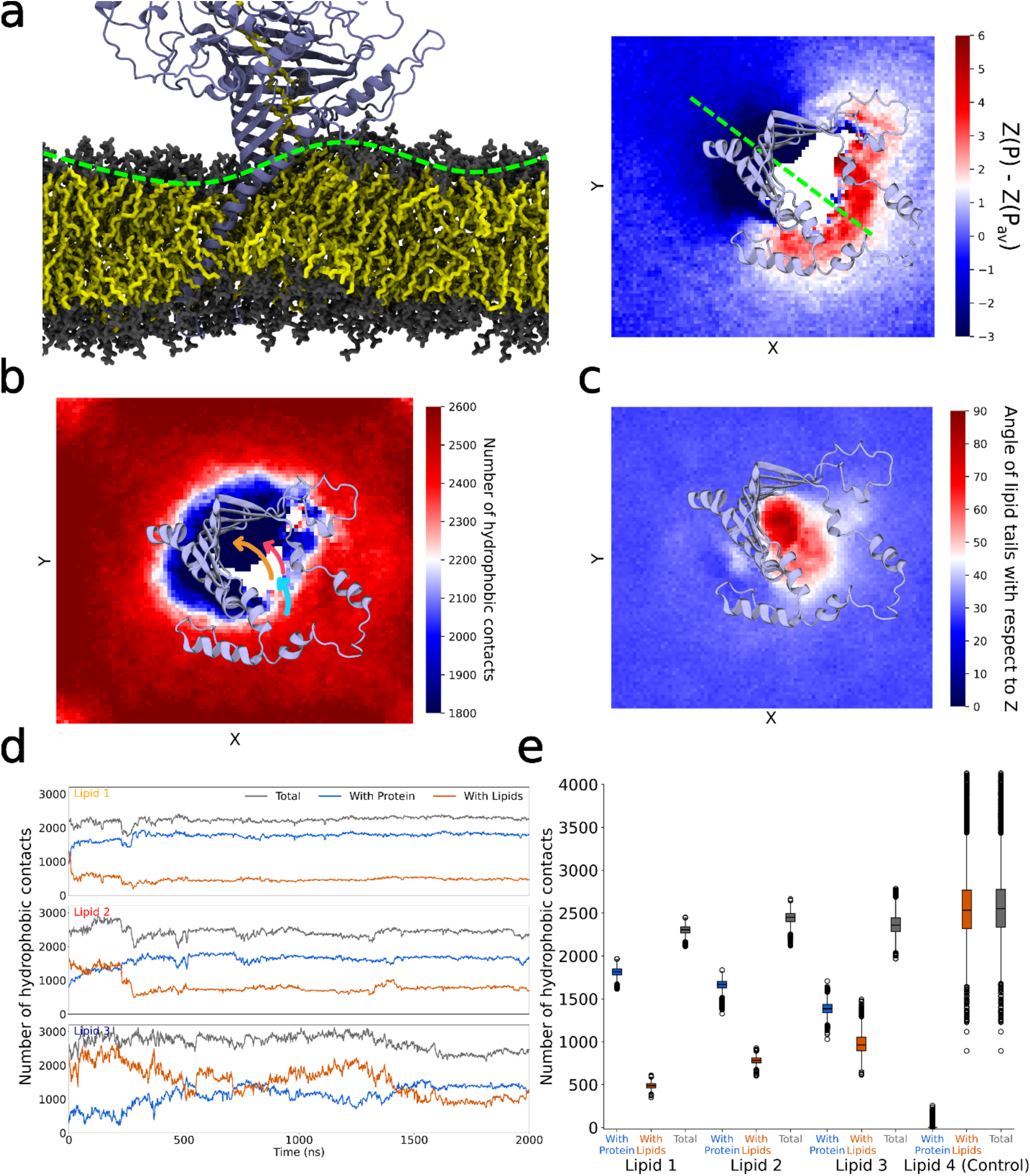
BLTPs destabilize membrane properties to promote lipid uptake. **a.** Snapshot of a simulation frame showing the disruption of the membrane close to the entry of the hydrophobic cavity (left), and *xy* map of the average difference in nm between the z values of the P atoms of the lipids with respect to the membrane centre (right), without considering the lipids that desorb. Red colour indicates positive curvature and a darker blue colour negative curvature **b.** *xy* map of the average number of lipid-lipid hydrophobic contacts, desorbed lipids are not considered. The arrows indicate the average movement of the center-of-mass of the desorbed lipids (same colors as in Fig. 1) during the entire trajectory **c.** *xy* map of the average angle of the lipid tails with respect to *z*, red colour corresponds to a perpendicular disposition, whereas a blue color is indicative of a parallel disposition with respect to *z.* **d, e.** Time traces **(d)** and boxplots **(e)**, respectively, of protein-lipid hydrophobic contacts (blue), lipid-lipid hydrophobic contacts (orange), and total number of hydrophobic contacts (grey) for the three lipids that are taken up by Csf1 (Lipids 1, 2 and 3), and a control lipid that remains in the membrane (Lipid 4). Boxplots were computed using the raw data from the last 500 ns of the trajectory in **(d)**. The results correspond to the simulation of the N-terminal part of Csf1 embedded in a DOPC bilayer

Interestingly, the shape of the N-terminal RBG domain of the protein creates a ‘C’ pattern of hydrophobic contacts within the lipid membrane (Fig. 3b, Fig. S4), with the lipids entering the hydrophobic cavity through the opening of the ‘C’, where lipid-lipid hydrophobic contacts are depleted. These observations hold true for all tested membrane compositions and proteins. As shown in the time traces of hydrophobic contacts for the desorbed lipids (Fig. 3d, Fig. S5), these lipids suffer a major loss of hydrophobic lipid-lipid interactions during the simulations (Fig. 3d, Fig. S5).

To test whether this loss of hydrophobic contacts with other lipids is compensated by the formation of new hydrophobic contacts with the protein, we computed the hydrophobic contacts between the desorbed lipids and all hydrophobic residues facing the interior of the Csf1 cavity. Indeed, when also considering these interactions, the number of hydrophobic interactions remains constant in most cases, with total numbers very similar to those of the lipids that remain in the membrane (Fig. 3d, e; Fig. S5).

Overall, our simulations indicate that BLTPs generate a favourable desorption path within their surroundings: the lipids close to the entry of the protein cavity undergo a loss of in-bilayer hydrophobic contacts (Fig. 3b, Fig. S3), which are only recovered when they either return to the bulk membrane, or enter the protein hydrophobic pocket, due to newly-formed hydrophobic interactions either with the protein non-polar residues or with the already-desorbed lipids that are present in the protein cavity (Fig. 3d,e, Fig. S5).

### The presence of BLTPs dramatically reduces the energy of lipid desorption

In the absence of BLTPs, lipid desorption from the lipid bilayer is energetically costly,^48,49^ and this step has been proposed to be rate-limiting for lipid transport by BLTPs.^38^ To quantify the energetic effect of the protein on the lipid desorption process, we computed the potential of mean force (PMF) using umbrella sampling simulations for DOPC lipids with and without the embedded BLTP. To this end, we compared the desorption energy profiles (Fig. 4a) of a DOPC lipid desorbing from a membrane without a BLTP (black line in Fig. 4b), and of the 3 lipids that fully desorb spontaneously inside the Csf1 cavity (orange, red, and blue lines in Fig. 4b) in our unbiased AA MD simulation (Fig. 1d, e).

**Figure 4.**
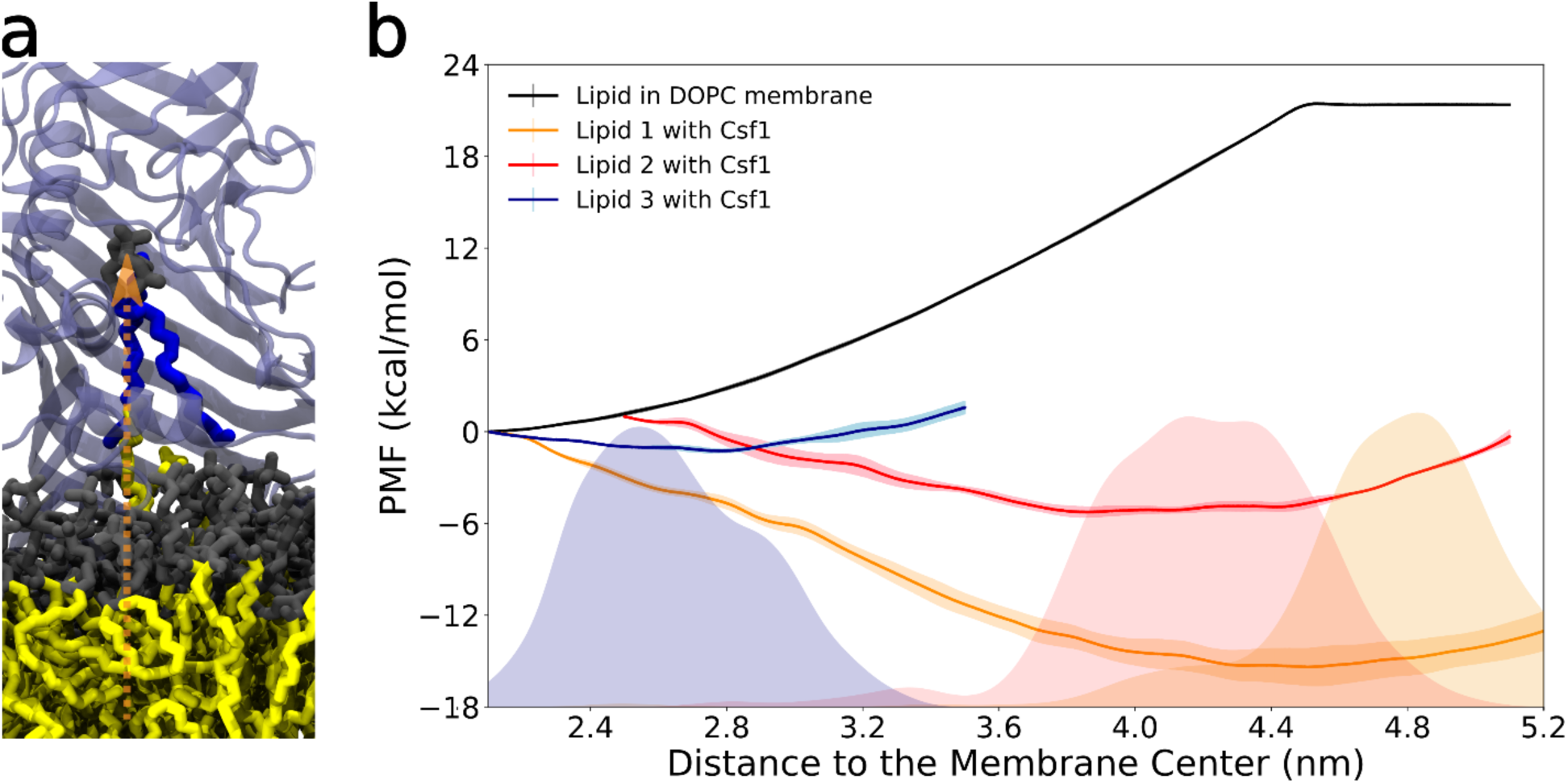
BLTPs promote the desorption of lipids. **a**. Schematic representation of the reaction coordinate: distance in z of the P atom of the desorbed lipid to the membrane center **b.** PMF profiles in kcal/mol, with errors, for the desorption of a lipid in a membrane without BLTP (grey line), and of the three lipids discussed in the unbiased simulations of Csf1 embedded in a DOPC membrane (see Fig. 1). In the background, the distance distribution plots from the corresponding unbiased AA MD simulation are displayed using the same color scheme.

The spontaneous desorption of lipids from pure lipid membranes has indeed a large energetic cost (Fig. 4b) and, therefore, is never observed during unbiased MD simulations. Using umbrella sampling, the computed energy barrier for the desorption of a lipid without the BLTP exceeds 20 kcal/mol, in agreement with previous reports,^50,51^ confirming the unfavourable nature of this process (Fig. 4b, black curve). On the other hand, when considering the desorption of individual lipids into the hydrophobic cavity of Csf1, the energy barrier for this process lowers dramatically, as the new environment for the lipid resembles the one within the membrane (Fig. 1f, Fig. 3e). In fact, as discussed previously, a stable number of hydrophobic contacts is only reached when the lipids simultaneously interact with the protein and with other desorbed lipid tails. Consequently, the desorption of the lipids in this scenario becomes energetically favourable, with minima in energy that correspond to the metastable states observed in our unbiased simulations (Fig. 4b). Specifically, the energy of desorption is of -15.2 kcal/mol for the first desorbed lipid, -6.1 kcal/mol for the second, and -1.3 kcal/mol for the third, highlighting the energetically favourable nature of the process in the presence of the BLTP.

Notably, the process of lipid desorption becomes less energetically favourable as the number of lipids inside the protein increases. On the one hand, this explains the low number of desorption events that we observe in our simulations: upon desorption of a limited number of lipids, the system reaches a metastable state in which the BLTPs are filled by the number of lipids that allows for the formation of the maximum number of hydrophobic contacts within the cavity, but that does not lead to an overcrowding of lipids that would result in steric clashes. On the other hand, our results indicate that removal of the desorbed lipids, e.g. via transport along the protein tunnel, either by active or passive mechanisms, could result in the spontaneous desorption of additional lipids and in the continuation of the process.

### Specific protein-membrane interactions are required for lipid uptake

Our simulations indicate that the interplay between the protein and the nearby membrane is crucial for lipid extraction and for the energetics of this process. To test whether altering interfacial protein-membrane interactions could affect lipid desorption, we focused on hydrophobic interactions between the inserted beta-sheets of the protein and the lipid bilayer. Specifically, we selected two highly conserved hydrophobic residues (Fig. 5a, b) located in the protein region extensively interacting with the headgroups of the lipids in the bilayer, and mutated them to charged ones (F80R, L99E). In our AA MD simulations, the presence of these mutations prevents the formation of stable hydrophobic interactions between the membrane lipids and the inserted protein beta-sheets, leading to a lack of desorption events (Fig. 5c). Concomitantly, the protein is not able to destabilize the bilayer, with only mild curvature (Fig. 5d) and no tilting of the lipid tails around the protein observed (Fig. 5e).

**Figure 5.**
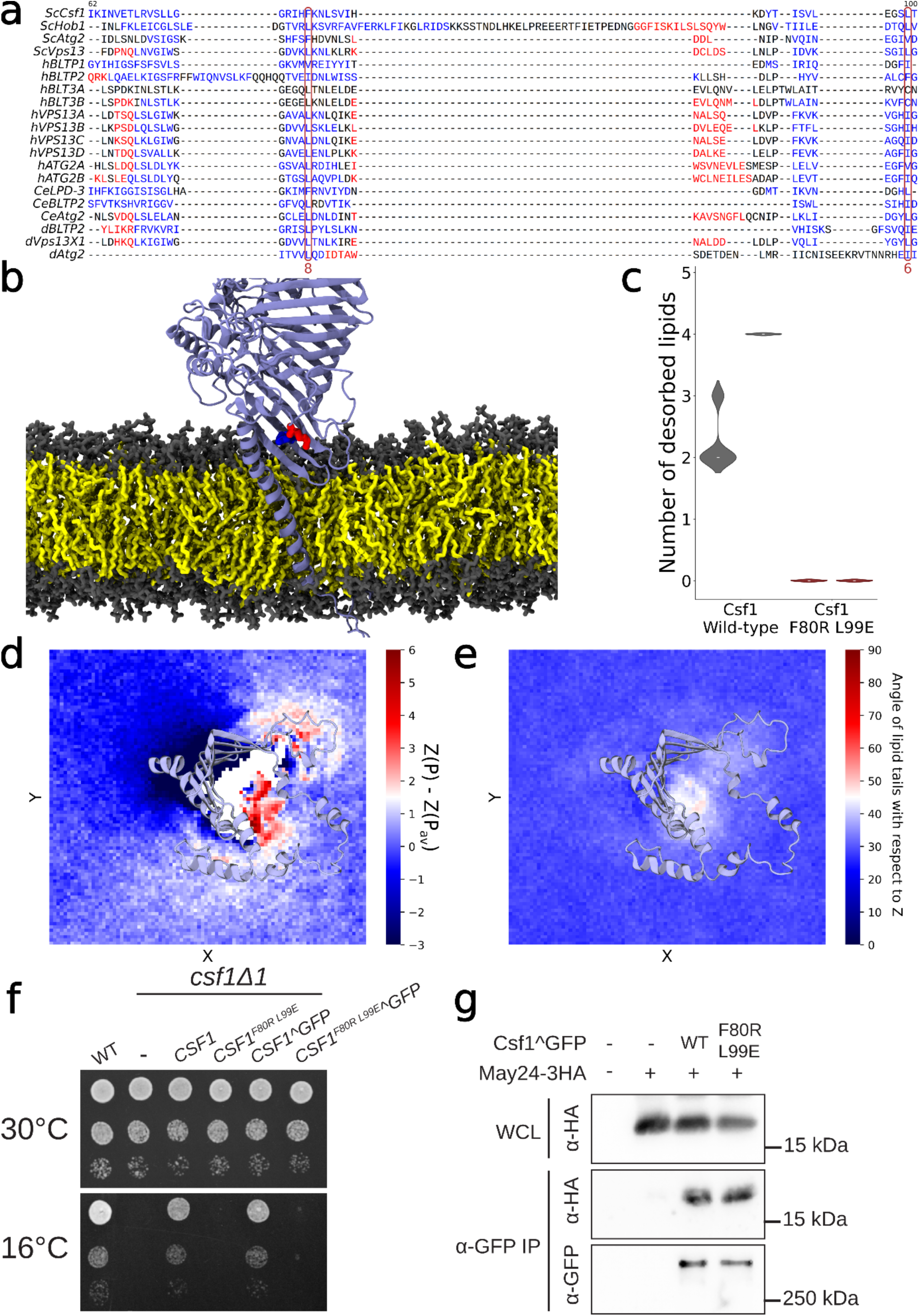
Disrupting Csf1’s interfacial interactions with the lipid bilayer blocks the desorption of lipids and impairs Csf1-mediated cold adaptation. **a.** Multisequence alignment of BLTPs across different organisms using PROMALS3D.^52^ The mutated residues F80, and L99 are shown in red boxes, both are conserved hydrophobic residues across all BLTPs. Conservation indexes of the mutated residues are displayed in red below the boxes.^53^ The color of the amino acids corresponds to the predicted secondary structure (red: alpha-helix, blue: beta-strand). **b.** Snapshot of the final frame of the simulation with the F80R, L99E double Csf1 mutant, no desorption of lipids is observed. Mutated residues are displayed in blue (F80R) and red (L99E). **c.** Violin plots with the number of desorbed lipids for the wild-type and the double mutant Csf1 simulations in DOPC bilayers for two different replicas. d. *xy* map with the average z values in nm of the P atoms of the lipids with respect to the membrane centre, red color is indicative of a positive curvature, whereas blue color is indicative of negative curvature; all lipids have been considered for the analysis. e. *xy* map with the average angle of the lipid tails with respect to *z*, red color corresponds to a perpendicular disposition whereas blue indicates a parallel disposition with respect to z **f**. Plasmid-expressed wild type Csf1, but not the Csf1^F80R L99E^ mutant, rescues the cold sensitive phenotype of the *csf1Δ1* null mutant in both untagged and GFP-tagged contexts. “-” indicates the empty vector control **g**. Immunoprecipitations of wildtype and mutant Csf1 show comparable protein levels and co-purification of the May24 lipid scramblase. “-” indicates expression of untagged proteins. WCL: whole cell lysate

To investigate if Csf1’s interaction with the lipid bilayer is indeed essential for its function in a biological context, we expressed the Csf1^F80R L99E^ mutant in a yeast strain lacking functional Csf1 (Fig. 5f). Previous studies have established that Csf1 is required for yeast cold adaptation, likely by maintaining membrane fluidity via its lipid transport activity.^54^ We found that expression of either untagged or internally tagged forms of wild type Csf1 fully rescued the cold-sensitive growth of the *csf1Δ1* mutant strain, whereas the Csf1^F80R L99E^ mutant was completely defective in either context (Fig 5f). To confirm that these mutations do not impact protein stability or interactions with other protein partners, we verified that the immunoprecipitated Csf1^F80R L99E^^GFP mutant was present at wild-type levels and co-purified with its known N-terminal interactor, the scramblase May24 (Fig. 5g).^45^ Taken together, this suggests that the Csf1 interfacial protein-membrane interactions are essential for function *in vivo*.

## Discussion

Recent observations about lipid transfer proteins are unveiling the key role of BLTPs in membrane and organelle homeostasis.^8,11,29^ Despite growing interest in this field, the details about how these proteins transfer lipids between membranes are still unresolved. Due to the inherent limitations of studying BLTPs experimentally, computational methods provide an important tool to increase our understanding of their structure and mechanisms. Our work, consisting of AA MD simulations of several BLTPs in membranes with different lipid compositions, confirms the ability of these proteins to sequentially take up multiple lipids. Additionally, our simulations reiterate that all BLTPs containing TM domains are capable of spontaneously desorbing multiple lipids without any apparent selectivity. Upon desorption, these lipids freely diffuse within the region of the hydrophobic cavity close to the membrane, changing conformation until reaching a metastable state.

Beyond confirming previous models of BLTPs’ mode of transport, our simulations provide novel molecular insights into the precise mechanism of lipid desorption and, ultimately, transport. First, our simulations indicate that lipid entry consistently takes place via the “C”-shaped opening of the RBG fold, with the lipid polar heads, which remain solvated during the desorption process, interacting with the polar residues at the entry of the cavity, while the acyl chains adopt a “splay” conformation and interact with hydrophobic residues inside the cavity.

Second, we found that this entry mechanism is promoted by the specific interactions that the BLTP establishes with the lipid bilayer: when embedded in lipid membranes, TM-anchored BLTPs alter bilayer properties by generating local curvature, by changing the orientation of the lipid tails in their proximity, and, most importantly, by depleting lipid-lipid hydrophobic interactions. Consequently, lipids located at the entry of the hydrophobic cavity are prone to undergo desorption to overcome the loss of such hydrophobic interactions. The ‘C’ shape of the protein facilitates this process by providing an amphipathic environment for the desorbed lipids: the headgroups remain solvated thanks to the opening of the ‘C’, whereas the lipid tails interact with both the protein hydrophobic residues as well as with the lipid tails of other desorbed lipids. Hence, the total number of hydrophobic contacts for the desorbed lipids remains almost constant, which explains the large number of desorption events observed in our simulations, regardless of the BLTP and the membrane composition. Interestingly, even if a comparable disruption of the membrane local hydrophobic environment has been proposed for shuttle LTPs, only very few or partial lipid desorption events have been observed for AA MD simulations of shuttle LTPs when compared to our simulations^38,48,55,56^.

To validate the relevance of membrane disruption for lipid desorption, we rationally designed a Csf1 double mutant with strongly-decreased membrane disruption activity, and we confirmed that this mutant has lost its ability to desorb lipids *in silico* as well as its physiological function *in vivo*. Unlike in previous studies,^32–36^ however, the designed mutations are not located within the hydrophobic channel. Rather, the simulations reveal an unexpected and essential role for conserved residues on the surface of the BLTP located at the interface with the bilayer, causing membrane curvature and disrupting lipid-lipid interactions. This observation opens further possibilities to interfere with the lipid-transfer ability of BLTPs, particularly since this mechanism appears to be conserved for all the proteins tested in this study (Fig. S3 and S4) and as the mutated residues appear to be conserved hydrophobic residues across BLTPs from different organisms (Fig. 5a).

Whether this mechanism remains valid for BLTPs that do not possess TM domains, such as Vps13/VPS13A-D and Atg2/ATG2A-B, requires further understanding of how these proteins attach to lipid membranes. Interestingly, both Vps13/VPS13A-D and Atg2/ATG2A-B have been proposed to interact with lipid bilayers through highly conserved *N*-terminal helical regions,^57–59^ which could play a similar membrane-disrupting role as what was observed here for TM-anchored BLTPs. In addition, how the interaction with other membrane-associated proteins, such as lipid scramblases^45,60–62^ could alter this process remains an active area of research.

Finally, by comparing the energy profiles for the desorption of lipids in the presence/absence of a BLTP, our simulations confirm that the presence of the protein completely abolishes the large energy barrier for this process, thus favouring the sequential desorption of multiple lipids into its hydrophobic cavity. As more lipids desorb into the protein, this process becomes energetically unfavourable due to the saturation of the hydrophobic residues inside the protein cavity and the formation of hydrophobic interactions with the already desorbed lipids, limiting the number of events observed in our simulations. These observations suggest that the flow of lipids towards the acceptor membrane would, in turn, spontaneously trigger more desorption events at the N-terminal part, facilitating continuous lipid transport. Taken together, our data suggests that BLTPs act as a natural “sink” for lipids, and that lipid desorption from the donor membrane is very unlikely to be the rate-limiting step for lipid transport.

## Methods

### System setup and simulation details

The structure of all proteins but LPD-3 was predicted using Alphafold2.^63^ The structure of LPD-3 is from PDB ID 9CAP.^24^ Missing loops in the structure were modelled using Modeller.^64^

The TM domain of the proteins was embedded in the corresponding lipid membrane using PPM.^65^ The CHARMM-GUI Membrane Builder^66^ was used to prepare the lipid bilayers with the compositions listed in Table 1. Each system was solvated with CHARMM TIP3P water and ionized with 0.15 M of NaCl. Simulations were performed using the GROMACS software,^67^ and the CHARMM36m force field.^68^ The standard minimization and equilibration protocol from CHARMM-GUI was used for each system, consisting in an energy minimization using the steepest descent algorithm followed by two NVT equilibrations and four NPT equilibrations where positional restraints on both the protein and the lipids were increasingly removed. The equilibration runs used the Berendsen thermostat and semi-isotropic barostat to maintain the temperature at 303.15 K and the pressure at 1 bar.^69^ The Nose-Hoover thermostat,^70^ with separate coupling for the protein, the membrane, and the ionized solvent, and a coupling time constant of 1.0 ps, was used to keep the system at 303.15 K during the production runs. The 1.0 bar pressure in the productions runs was maintained with a semi-isotropic Parrinello-Rahman barostat,^71^ with a compressibility of 4.5·10^-5^ bar, a coupling time constant of 5.0 ps, and a 10-step frequency for the coupling. The Particle Mesh Ewald (PME) method was used to compute the electrostatic interactions, with a Fourier spacing of 0.12 nm and a cutoff of 1.2 nm.^72^ Van der Waals interactions were switched to zero over the 1-1.2 nm range. The LINCS algorithm was used to constrain the bonds involving hydrogen atoms. Periodic boundary conditions were employed in all three directions. Frames were written every 100 ps, with a time step of 2 fs. VMD was used for visualization and rendering.^73^

### Umbrella sampling simulations

The distance of the P atom of the desorbed lipid along the bilayer normal (*z*) with respect to the membrane center, which was calculated using the P atoms of the remaining lipids, was used as the reaction coordinate for the umbrella sampling simulations. The windows were spaced with a distance of 0.1 nm, and the total number of windows for each system is reflected in Table S1. Each window was simulated for 205 ns, using a harmonic force constant of 500 kJ·mol^-1^·nm^-2^. The first 5 ns were not considered, and the PMF was constructed using WHAM as implemented by GROMACS.^67^

In the case of the systems with the protein, the initial structure for each window was taken from unbiased MD simulations. In the system without the protein, the windows were selected from a pulling simulation of the P atom with respect to the membrane center, using a force constant of 500 kJ·mol^-1^·nm^-2^ and a pulling rate of 0.05 nm·ns^-1^.

### Analyses

Computation of all distances and angles was performed using the gmx distance and gmx gangle tools, respectively. In the case of the distance in *z* to the membrane center, the center was computed as the center of mass of all the P atoms of the membrane bilayer, excluding the ones of the lipids that desorb throughout the simulation. The distance was determined for the P atoms and for each of the terminal carbon atoms of the lipid tails (LT1 corresponds to C2X and LT2 to C3Y). A lipid was counted as desorbed when the distance between its P atom and the membrane center was greater than 3 nm. For the calculation of the angle with respect to *z*, the two vectors formed by the P atoms and the terminal carbons of the lipid tails were considered.

An in-house tcl script was used to calculate the solvation number of the lipid tails and headgroups, counting the number of water molecules within 5.0 Å of the corresponding set of atoms, excluding the hydrogen atoms.

The grid maps of curvature, angles of the lipid tails, and hydrophobic contacts were computed using python scripts with the libraries MDAnalysis,^74^ NumPy,^75^ Multiprocessing, and Pandas,^76^ taking into account only the lipids that are located in the leaflet where the desorption occurs. Before doing the analysis, the trajectory was fitted in the *xy* plane to the backbone atoms of the protein residues located at the interface with the membrane (43-143 for Csf1, 48-151 for BLTP1, 32-157 for Hob1, and 48-162 for LPD-3) using gmx trjconv with the -fit rotxy transxy option. The results of all trajectories were averaged within ranges of 1.0 Å in the *xy* plane.

For the curvature, the coordinates of the P atoms in each frame were printed to a separate file and then the curvature was estimated by calculating the difference in the *z* coordinate between all the individual P atoms of the lipids and the average *z* value of these atoms.

Regarding the grid map of the angles, the angle of the normalized vectors formed by the P atoms and the terminal carbon atoms of the lipid tails with respect to *z* were printed to a separate file along with the *x,* and *y* coordinates of the terminal carbon atoms.

The number of hydrophobic contacts was calculated using a similar approach as in ^38^, counting all the pairs of hydrophobic carbons within less than 1 nm. The number of pairs for each lipid and frame was added and printed to a separate file along with the *x*, and *y* coordinates of the center of mass of the corresponding lipid. The hydrophobic carbons of all phospholipids include atoms C23-C2X and C33-C3Y, whereas the hydrophobic carbons of the ergosterol molecules include the C1, C2 and C4-C28 atoms. A similar approach was used to count the number of hydrophobic contacts between the lipids and the protein. For all the proteins, the hydrophobic carbons include the side chain carbons of Ala, Ile, Leu, Met, Phe, Trp, Val, Pro, and Cys residues.

### Yeast Strains and Plasmids

Plasmids, primers, and yeast strains are outlined in Supplemental Table S2, S3, S4; all strains, plasmids and sequences are available on request. Homologous recombination-based integration of PCR products^77,78^ was used for gene deletions, which were confirmed by PCR of genomic DNA. To create a *csf1* loss of function mutation that does not perturb expression of the neighboring *GAA1* gene (*csf1Δ1*), codons for residues 1003-2018 were replaced by a NATMX4 selectable marker.

Plasmids were constructed in *Escherichia coli* via Goldengate assembly^79^ and confirmed by whole plasmid sequencing (Plasmidosaurus or Flow Genomics). To create centromere-based Goldengate yeast expression entry vectors, the following parts plasmids from the MoClo-YTK kit^80^ (Addgene Kit #1000000061) were assembled: pYTK008 (1 CON LS’), pYTK073 (5 CON RE’), pYTK081 (7 CEN6/ARS4), pYTK083 (8 AmpR-ColE1), pYTK047 (234 GFPdropout) and either pYTK074 6 URA (to make pMD628, which carries the *URA3* marker), pYTK075 6 LEU (to make pMD627, which carries the *LEU2* marker), or pYTK079 6 Hph (to make pMD626, which carries the HygB marker).

*CSF1* and *MAY24* plasmids were then made by Goldengate assembly using multiple fragments from which internal BsaI sites were removed using silent mutations and joined using MoClo-YTK syntax. To create a wild type *CSF1* clone, PCR was used to generate 4 successive fragments of *CSF1* from BY4741 genomic DNA using primers listed in Table S4, which were assembled with the *SAC6* promoter from pYTK022^80^ into pMD626 (to make pMD788) and pMD627 (to make pMD763).

To create *CSF1* plasmids with an internal GFP-Envy tag, the first *CSF1* fragment was instead amplified from genomic DNA of yeast strain LCY5726, which has a GFP Envy tag^81^ inserted after amino acid residue 806,^45^ then assembled as above to create pMD789 and pMD764. To assemble plasmids containing F80R and L99E mutations, the first *CSF1* fragment was replaced with a 449bp gene synthesis fragment encoding the mutations (TWIST Biosciences) and another fragment with the remaining sequence amplified using primers LC7664 and LC7644 from genomic DNA of either LCY5726 (for mutant *csf1* with internal ENVY tag, to create pMD791 and pMD722) or BY4741 (for mutant *csf1* with no tag, to create pMD790).

*MAY24* expression plasmids were similarly constructed by Goldengate assembly. In brief, the *MAY24* promoter was amplified from yeast genomic DNA with oligos LC7439 and LC7440. *MAY24* coding sequences were amplified with oligos LC7169 and LC7249 for untagged *MAY24*, or with LC7169 and LC7170 for assembly with a C-terminal tag. The untagged *MAY24* plasmid (pGS14) was made by assembly of the *MAY24* promoter and coding fragments, pYTK052 SSA1t, and pMD628. The *MAY24-HA* plasmid (pGS15) was assembled in a similar manner, but with addition of pMD712 3HA 3b parts plasmid.

### Growth Assays

Liquid cultures were grown overnight in complete yeast peptone dextrose media (YPD) (1% [w/v] yeast extract, 2% [w/v] Bacto-Peptone, 2% [w/v] glucose, 0.032% [w/v] tryptophan) containing 200 μg/mL hygromycin B (Cedarlane) and diluted to a concentration of 0.2 OD^600^/mL. 10-fold serial dilutions were plated onto two sets of YPD + hygromycin B plates and incubated at 30°C for 24 hours, or at 16°C for 4 days. Images were processed using Adobe Photoshop 2023.

### Co-immunoprecipitations

40 OD_600_/mL of plasmid-transformed yeast (GSY44) cells were grown to log phase, harvested, converted to spheroplasts by breakdown of cell walls with the Zymolase enzyme (SK1204911; MJS BioLynx) and stored at -80 °C. Spheroplasts were lysed in 500μl lysis buffer (50 mM HEPES pH 7.4, 1% Nonaethylene glycol monododecyl ether [C12E9], 50 mM NaCl, 1 mM EDTA, 1 mM PMSF, and 1X yeast/fungal Protease Arrest). Laemmli Sample Buffer (50 mM Tris-Cl pH 6.8, 2% SDS, 10% glycerol, 0.001% bromophenol blue, 2% β-mercaptoethanol) was added from a 2x stock solution to 50μl of lysate. Polyclonal rabbit anti-GFP (EU2; Eusera) was added to 430 μl of the remaining lysate and incubated for 1 hr at 4°C, followed by a second 1 hr incubation at 4°C with Protein A-Sepharose beads (10-1141; Thermo Fisher Scientific). Washed beads were resuspended in 50μl of Thorner Buffer (40mM Tris-Cl pH 6.8, 0.1 M EDTA, 5% SDS, 8M Urea, 0.04% bromophenol blue), and proteins were eluted from the beads by incubating samples at 70°C for 5 min.

After separation on SDS-PAGE gels, proteins were transferred to nitrocellulose membranes (50-206-3328; Fisher Scientific) and blotted with either mouse monoclonal α-HA.11 (clone 16B12, MMS-101R; Covance; RRID:AB_291262) or α-GFP (clones 7.1 and 13.1, 11814460001; Roche; RRID:AB_390913) antibodies. Horseradish peroxidase-conjugated polyclonal goat α-mouse (115–035-146; Jackson ImmunoResearch Laboratories; RRID: AB_2307392) was added as a secondary antibody. Blots were developed with ECL Prime chemiluminescent reagents (RPN2232, Cytiva) and imaged on an VILBER Fusion FX imaging system. Images were processed using ImageJ and Adobe Photoshop 2023.

## Acknowledgments

We thank Karin Reinisch for comments regarding this manuscript, as well as lab members from the Vanni lab. The MoClo-YTK plasmid kit was a gift from John Dueber. We gratefully acknowledge support from the Swiss National Science Foundation (grants CRSII5_189996 to SV), and the European Research Council under the European Union’s Horizon 2020 research and innovation program (grant agreement no. 803952, to SV), the Canadian Institutes of Health Research (grants OGB-177941 and PJT-180544 to EC) and a 2020 Canada Foundation for Innovation award (Innovation Funds Project 39914). This work was supported by grants from the Swiss National Supercomputing Centre under projects ID s1269, lp24 and lp69. DAL acknowledges support from the Margarita Salas program 2021–2023 funded by Ministerio de Universidades (MU-21-UP2021-030-53773022), while PCB was supported by a BC Children’s Hospital Research Masters studentship.

## Supplementary Information

### Supplementary Data

**Table S1.**
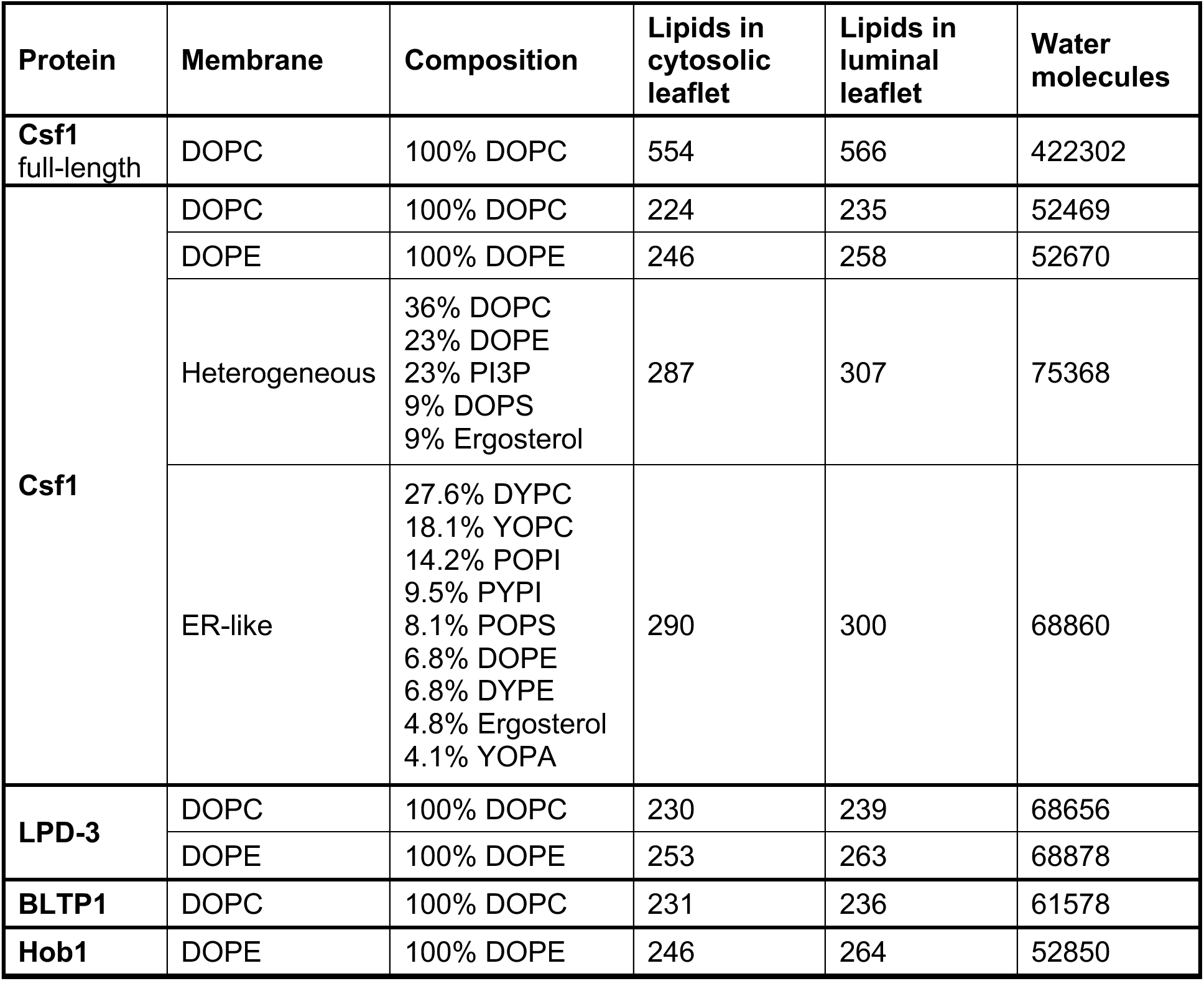
Composition (in %) of each lipid type for all the protein-membrane systems simulated, with the total number of lipids per leaflet and the number of water molecules in the simulation box.

**Table S2.**
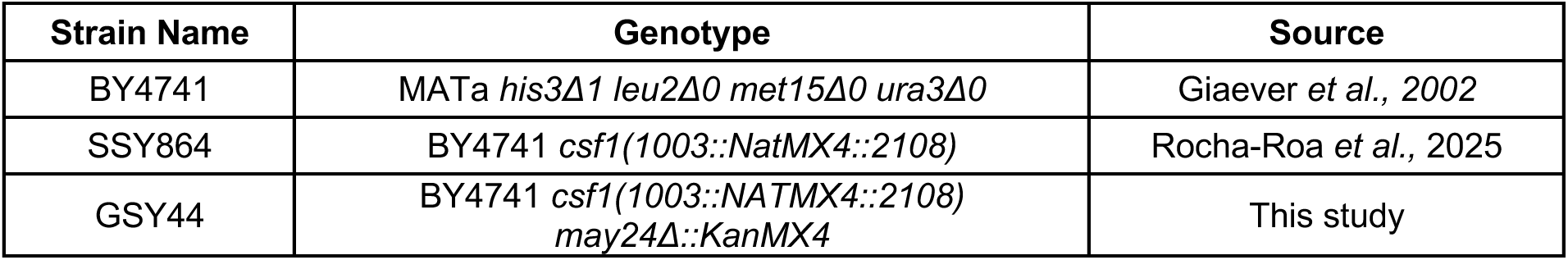
Yeast strains used in this study.

**Table S3.** Yeast plasmids used in this study.

| Plasmid Name | Description | Source |
| --- | --- | --- |
| pFA6-kanMX4 | pFA6-kanMX4 | Pringle lab (Longtine *et al.*, 1998) |
| pNR35 | pRS CEN Hph | This Study |
| pYTK022 | pSAC6pr Type 2 Parts plasmid | Gift from John Dueber (AddGene (#1000000061)) |
| pYTK052 | pSSA1t Type 4 parts plasmid | Gift from John Dueber (AddGene (#1000000061)) |
| pMD626 | pCEN Hph Goldengate entry vector | This Study |
| pMD627 | pCEN LEU Goldengate entry vector | This Study |
| pMD628 | pCEN URA Goldengate entry vector | (Rocha-Roa *et al.*, 2025) |
| pMD712 | p3HA 3b parts plasmid | This Study |
| pGS14 | pNativepr-May24(no-intron)-SSA1t (URA, CEN) | This Study |
| pGS15 | pNativepr-May24(no intron)-3HA-SSA1t (URA, CEN) | This Study |
| pMD763 | pSac6pr-CSF1 (LEU, CEN) | This Study |
| pMD764 | pSac6pr-CSF1<sup>806</sup>ENVY (LEU, CEN) | This Study |
| pMD772 | pSac6pr-CSF1 (F80R/L99E)<sup>806</sup>ENVY (LEU, CEN) | This Study |
| pMD788 | pSac6pr-CSF1 (HPH, CEN) | This Study |
| pMD789 | pSac6pr-CSF1<sup>806</sup> ENVY (HPH, CEN) | This Study |
| pMD790 | pSac6pr-CSF1 (F80R/L99E) (HPH, CEN) | This Study |
| pMD791 | pSac6pr-CSF1 (F80R/L99E)<sup>806</sup> ENVY (HPH, CEN) | This Study |

**Table S4.**
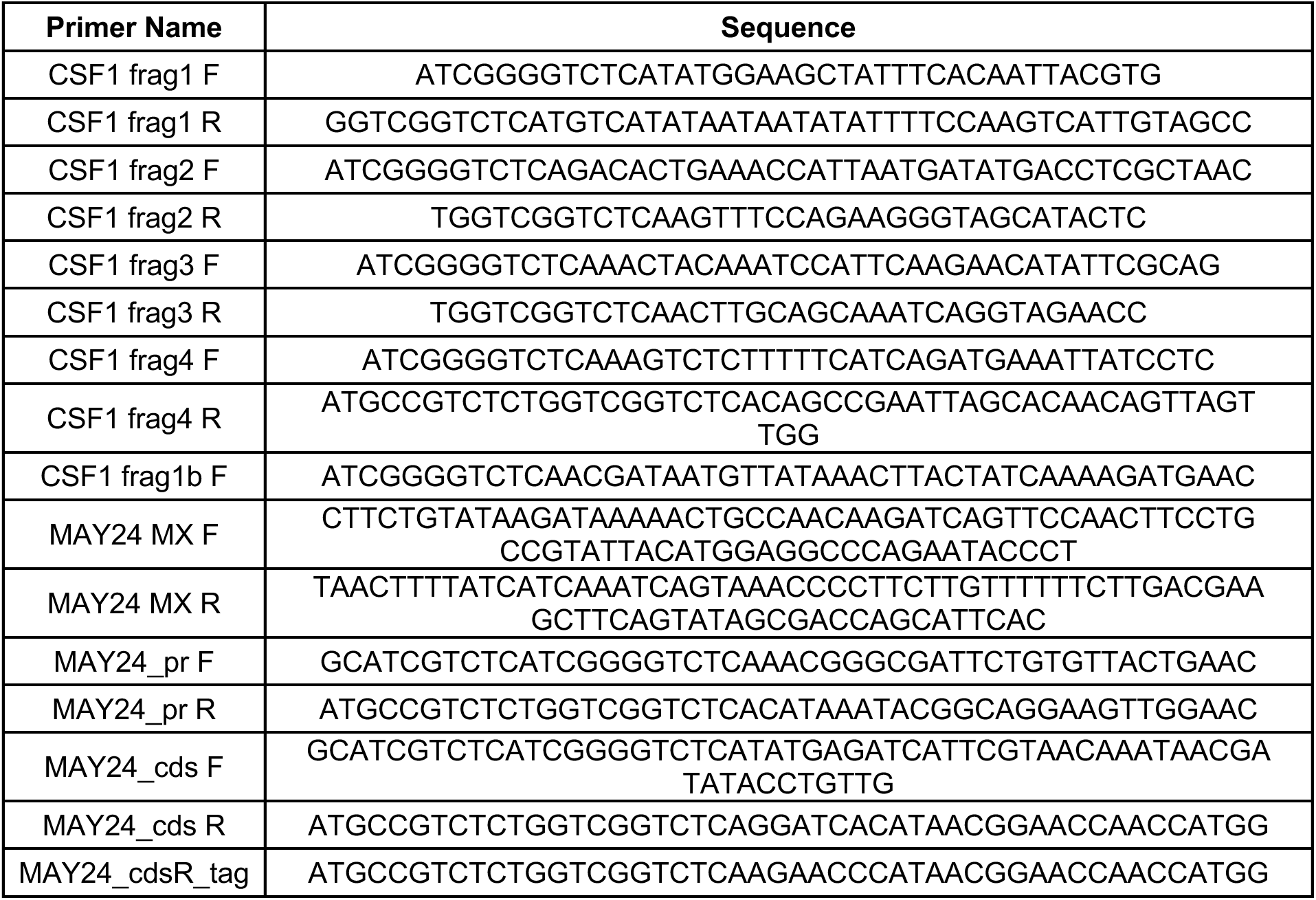
Yeast primers used in this study.

**Figure S1.**
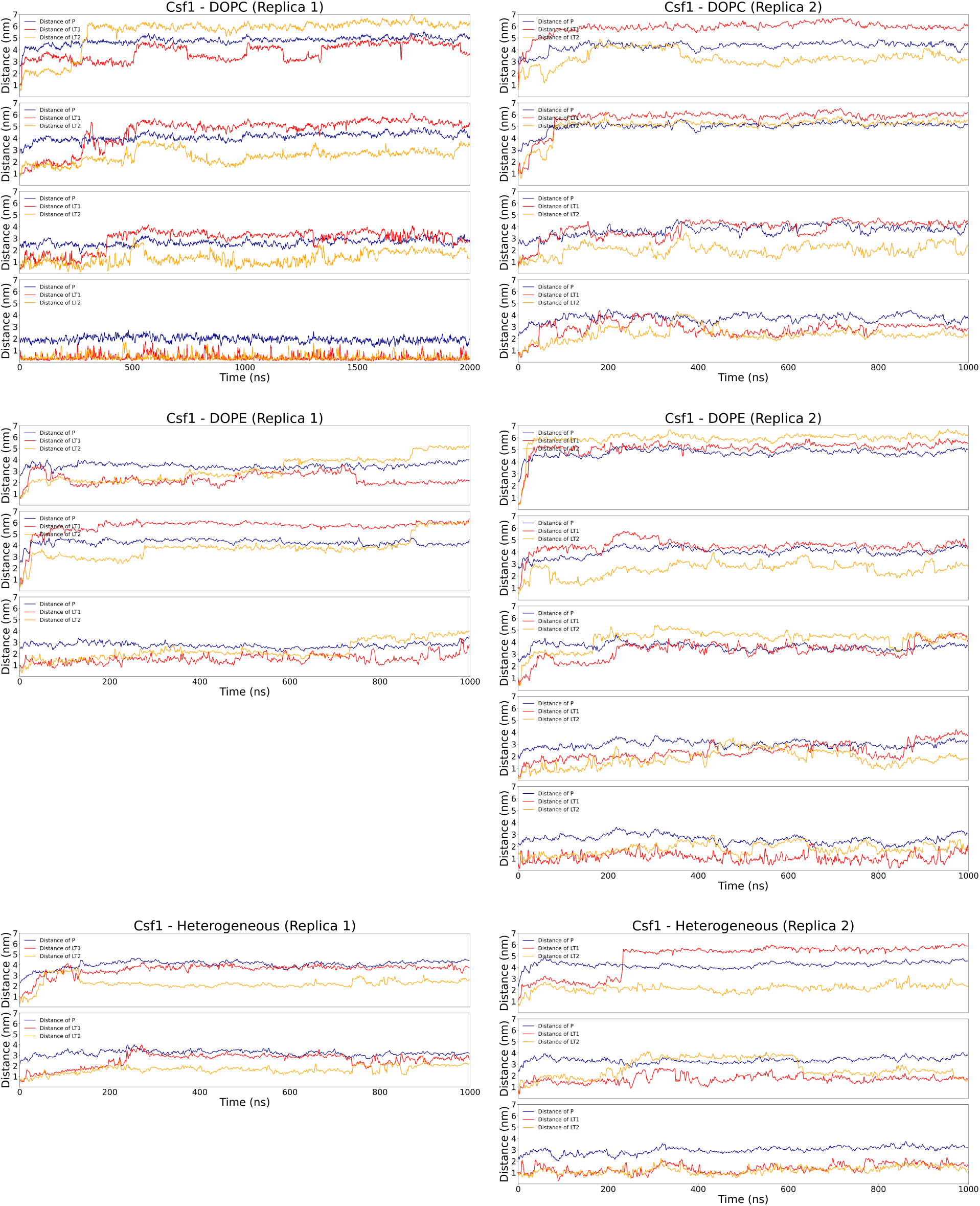
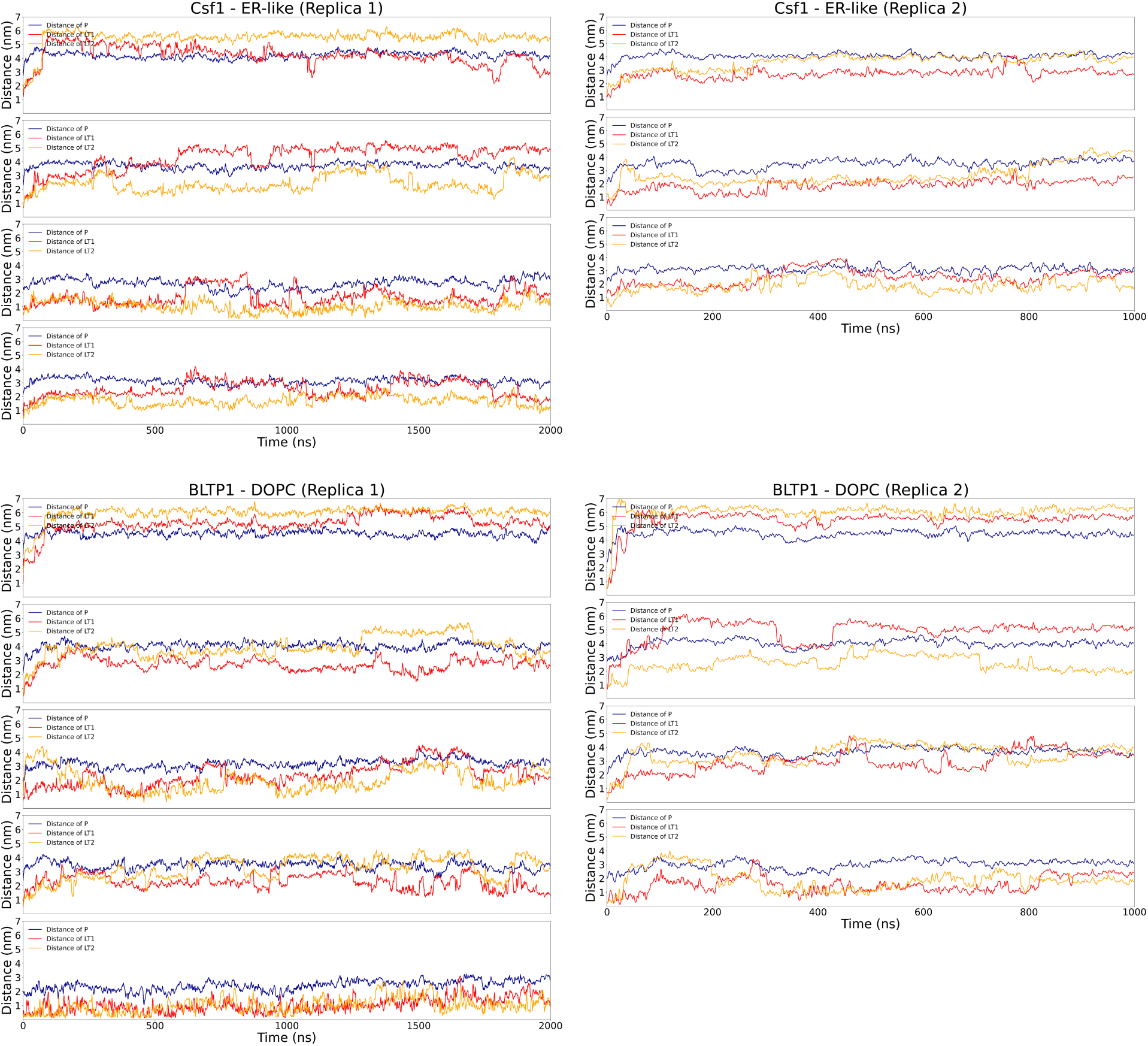
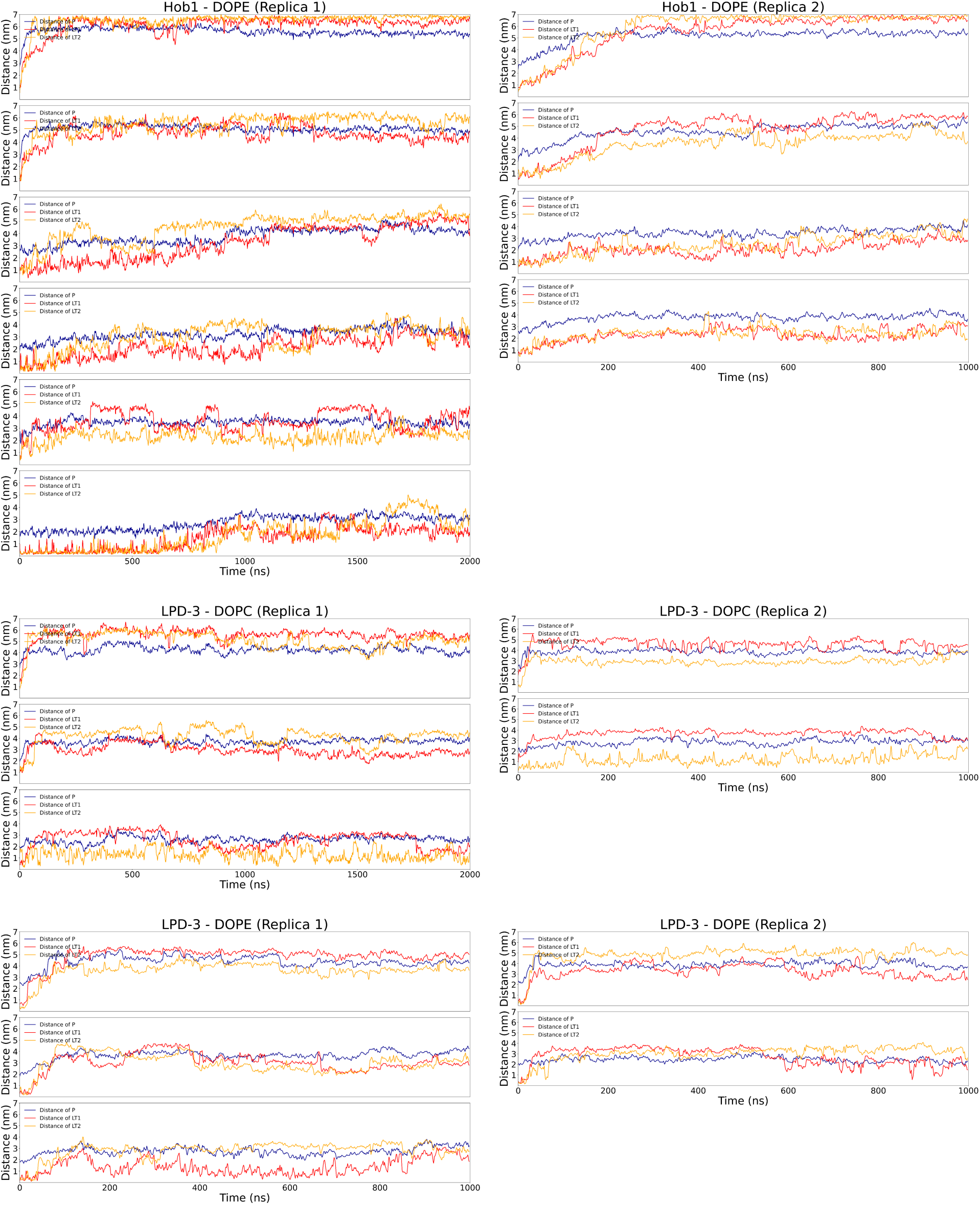
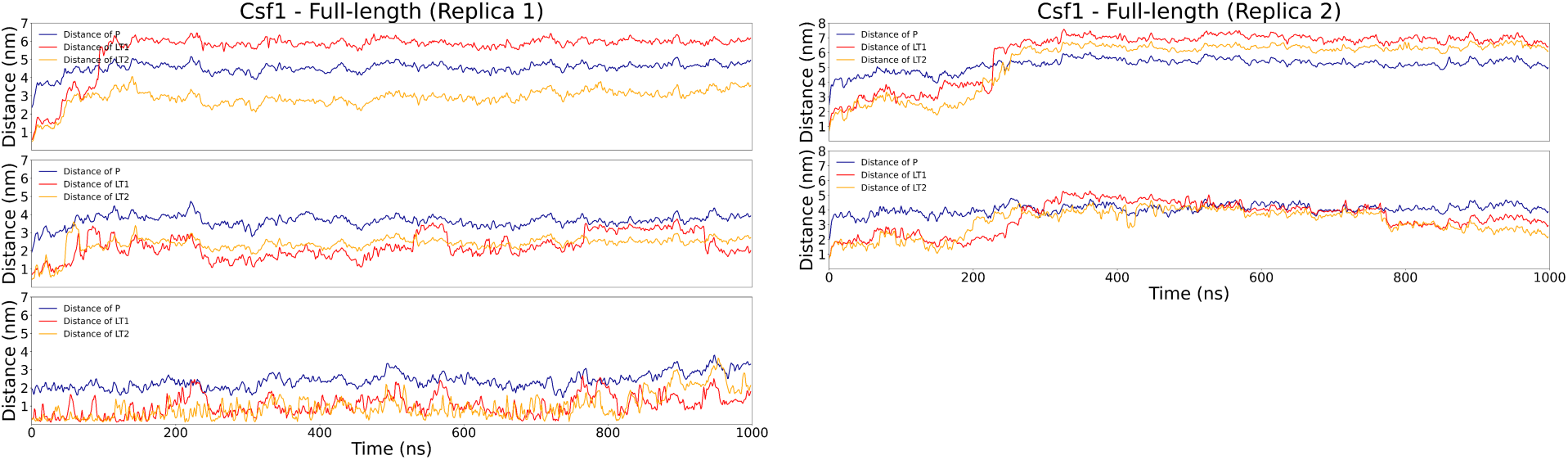
Time traces of the distance in *z* from the membrane center of the P atoms (blue), and terminal carbon atoms of the lipid tails (LT1, red, corresponds to C2X, LT2, orange, corresponds to C3Y) of desorbed or snorkelling lipids in all simulations. All the plots have been smoothened using block averages every 10 frames.

**Figure S2.**
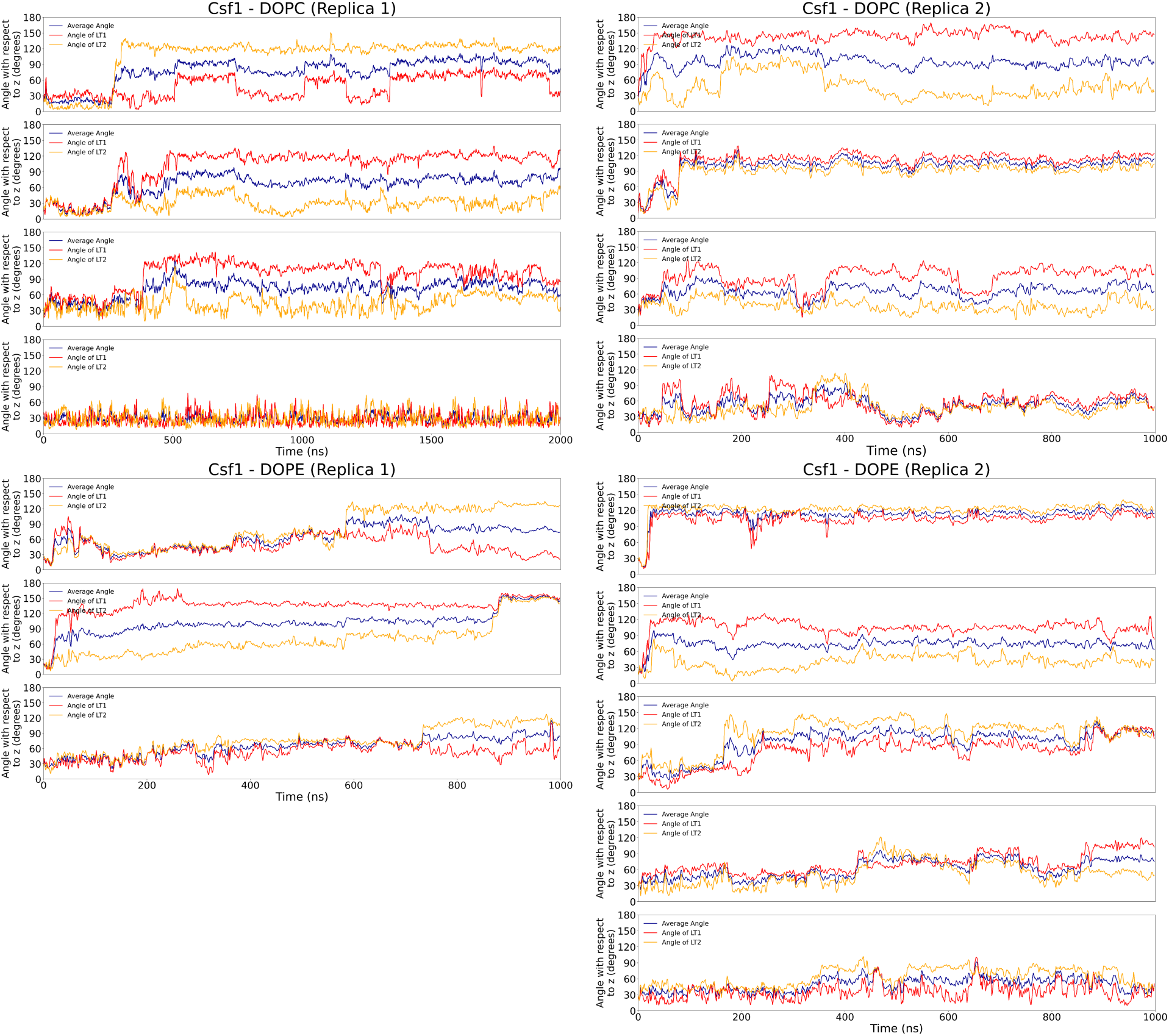
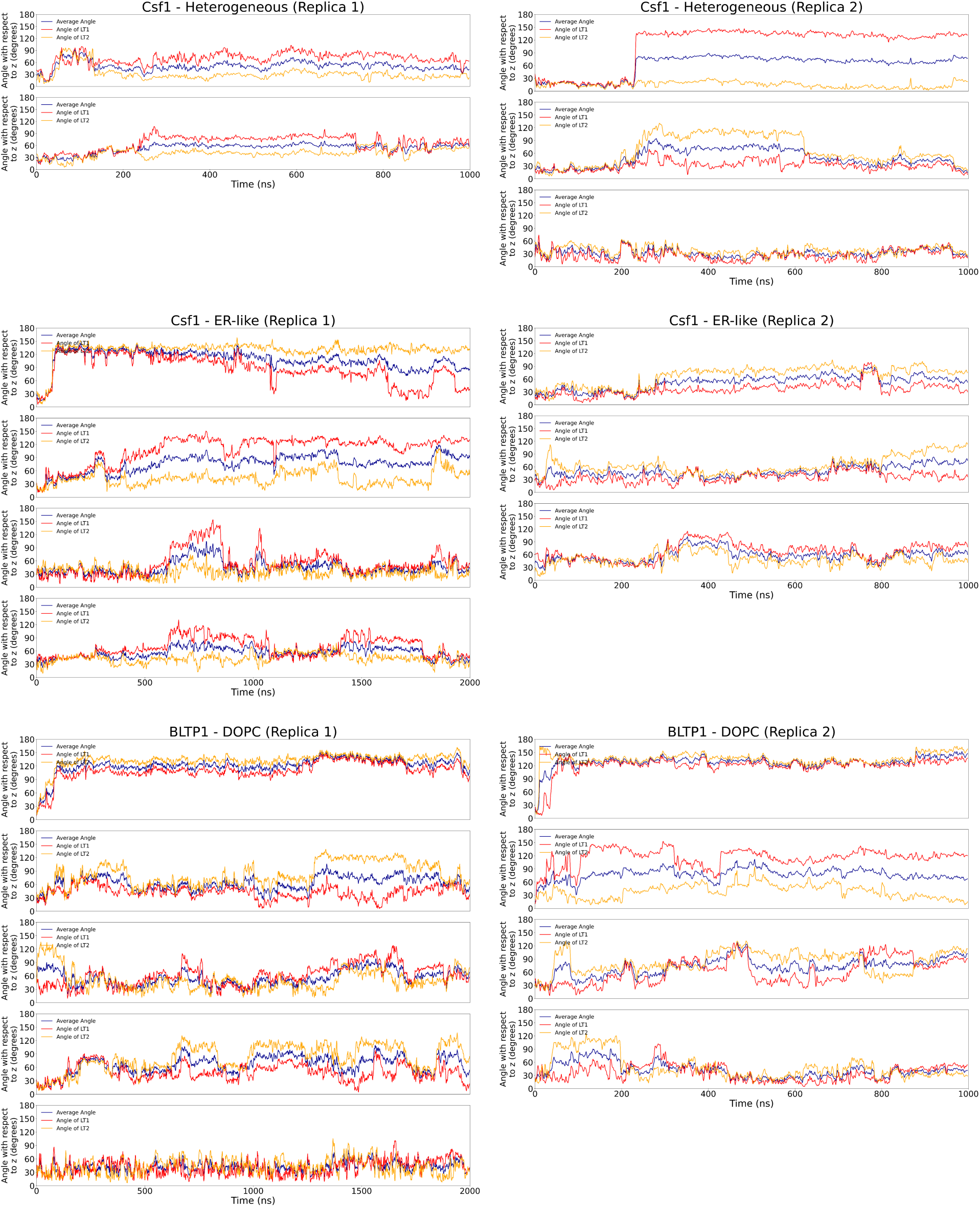
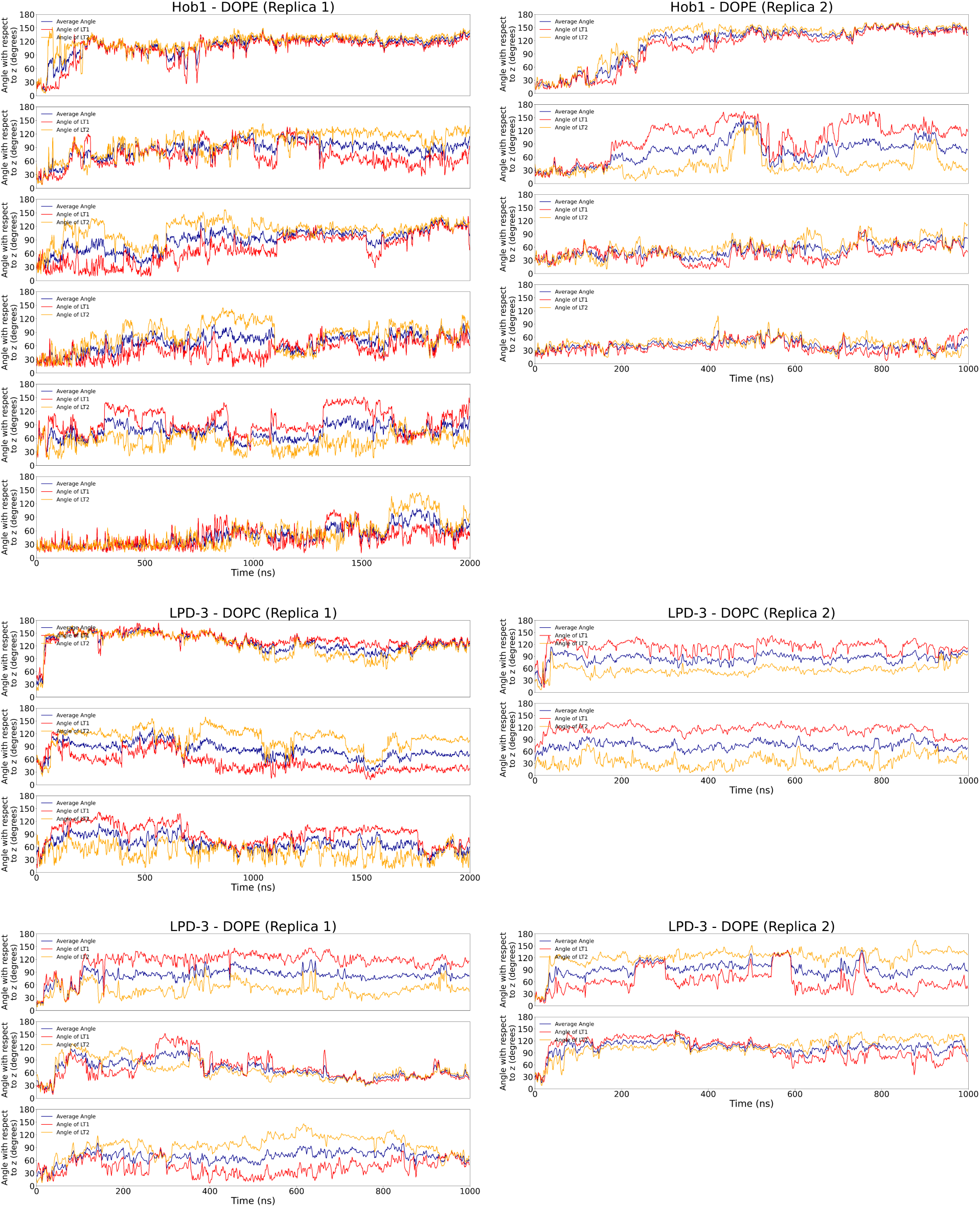
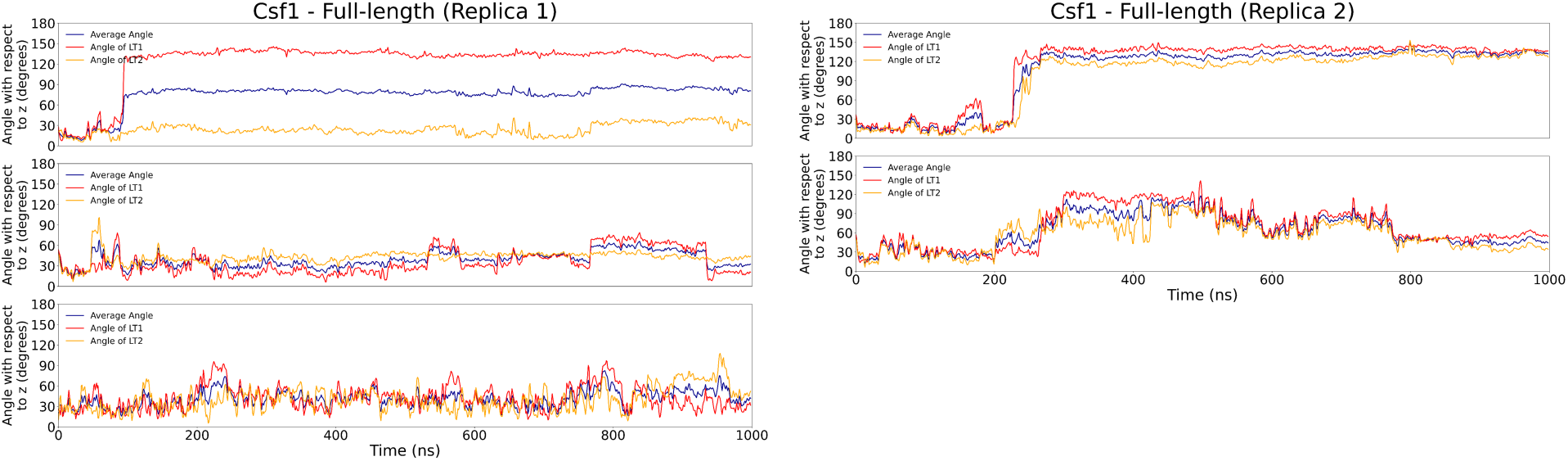
Time traces of the angles of the vectors formed by the P atoms and the terminal carbon atoms of the lipid tails (LT1, red, corresponds to C2X, LT2, orange, corresponds to C3Y) with respect to *z*. The average value of both angles is depicted in blue. All the plots have been smoothened using block averages every 10 frames.

**Figure S3.**
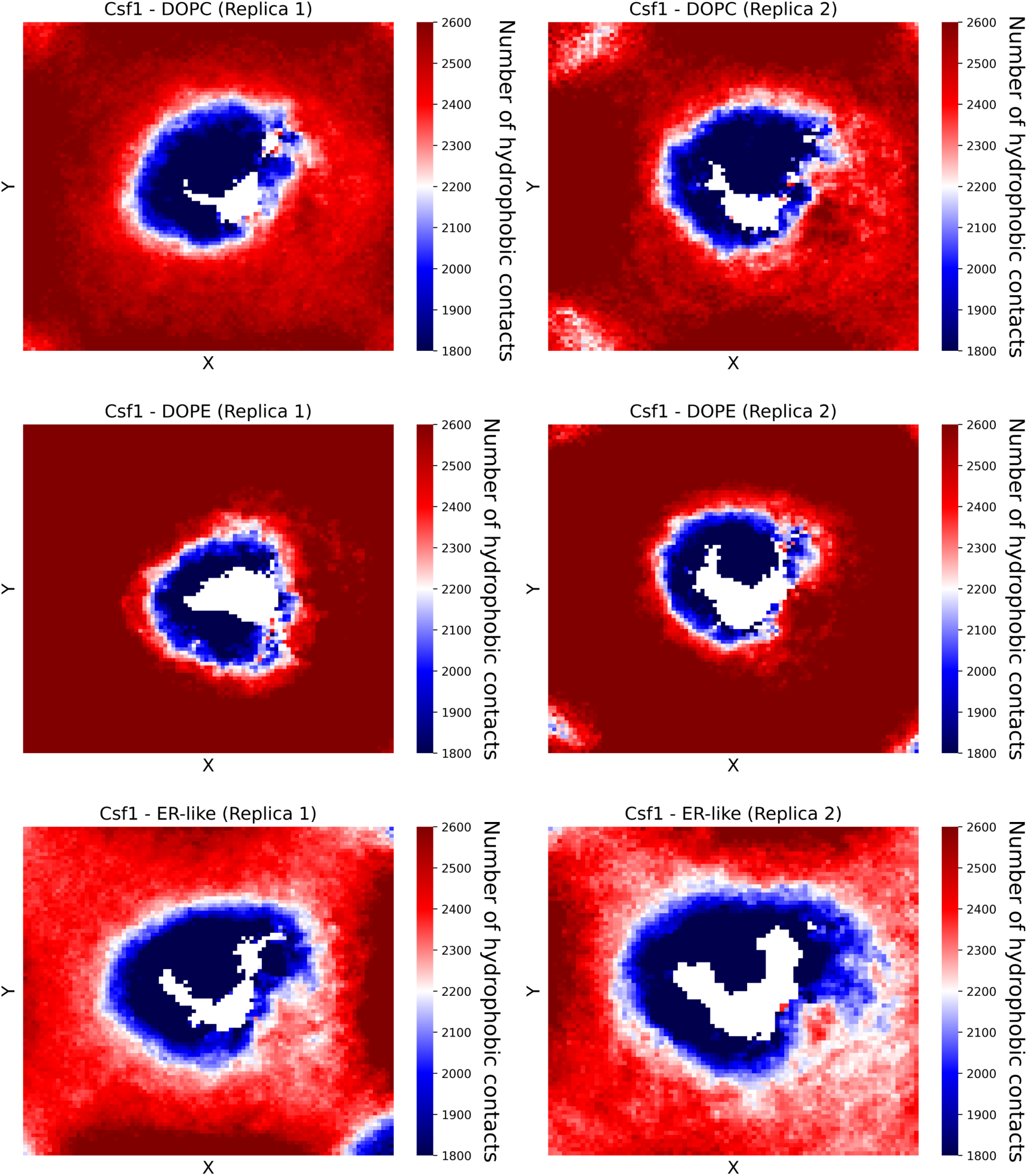
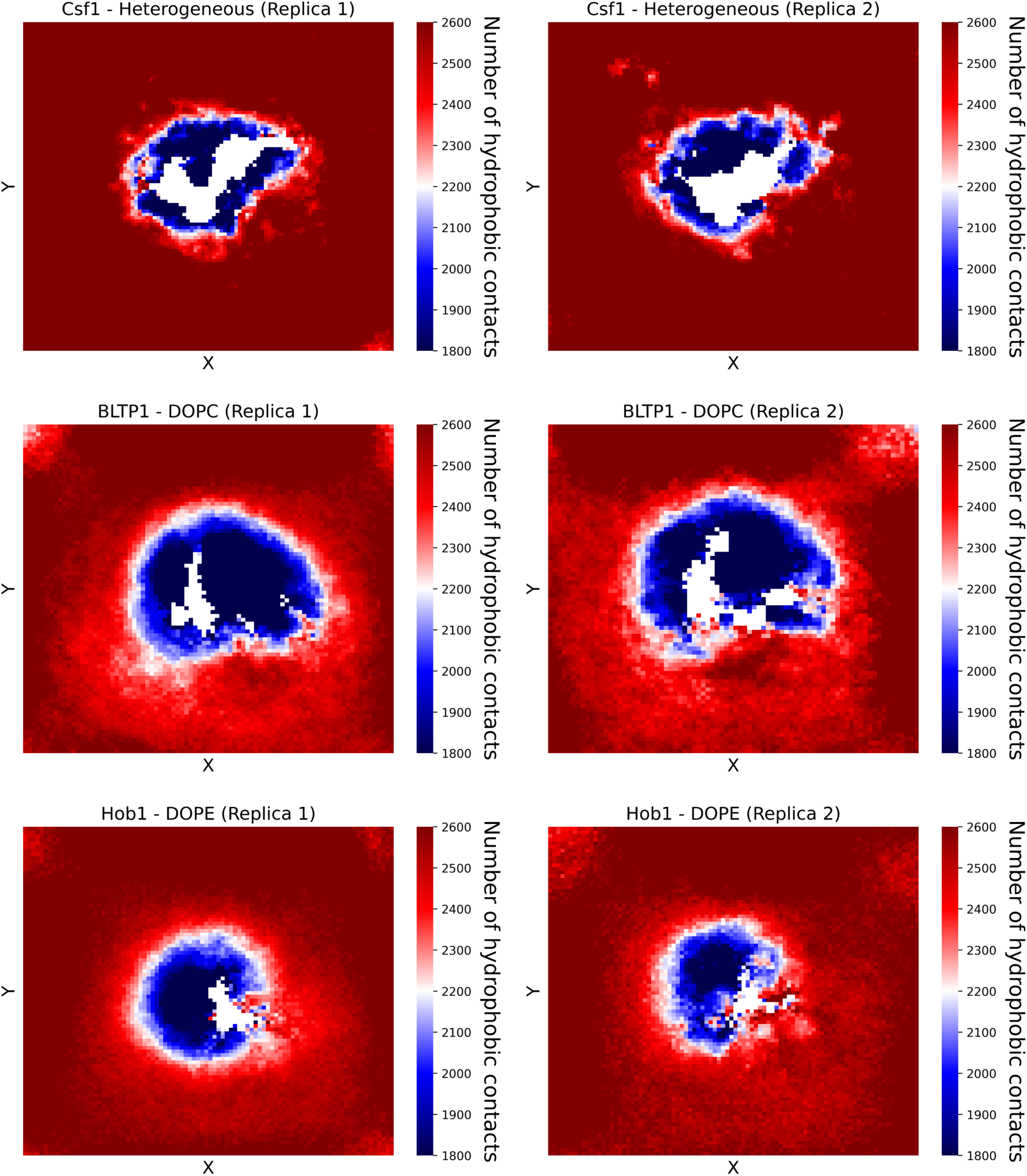
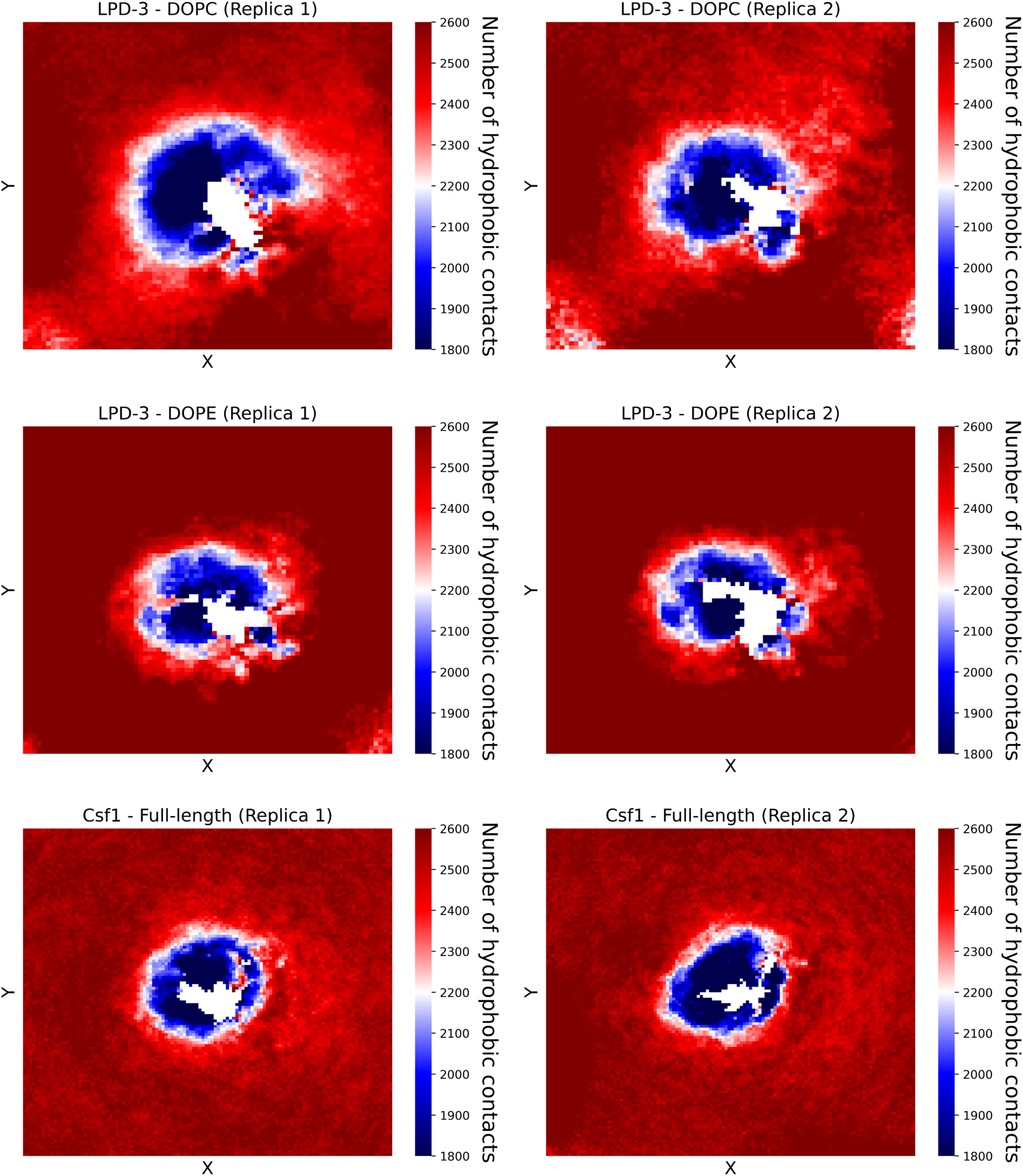
Grid *xy* maps of the average number of lipid-lipid hydrophobic contacts, without considering the lipids that desorb throughout the simulations. A red colour is indicative of a highly hydrophobic environment, whereas a dark blue colour corresponds to a low number of hydrophobic contacts.

**Figure S4.**
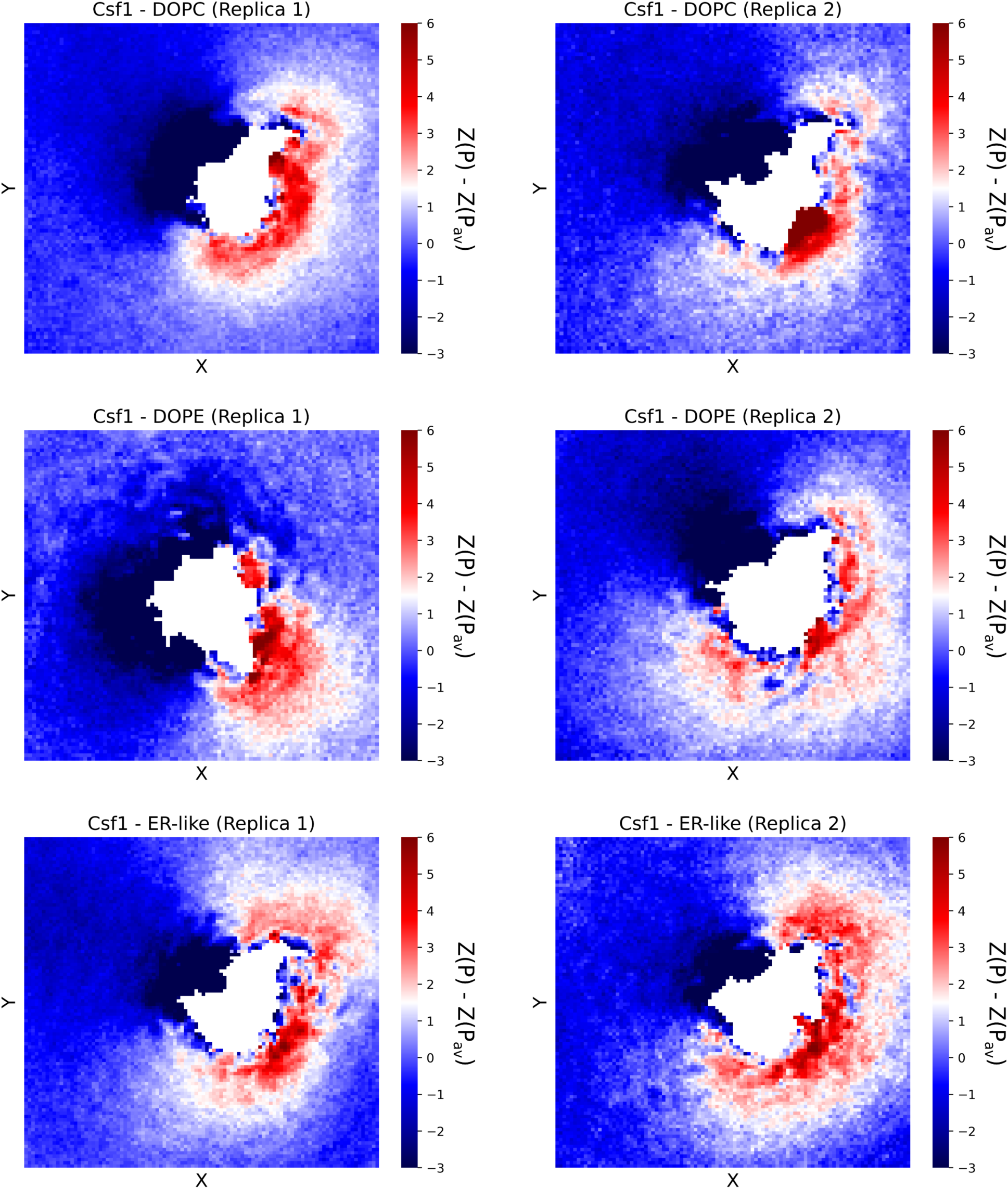
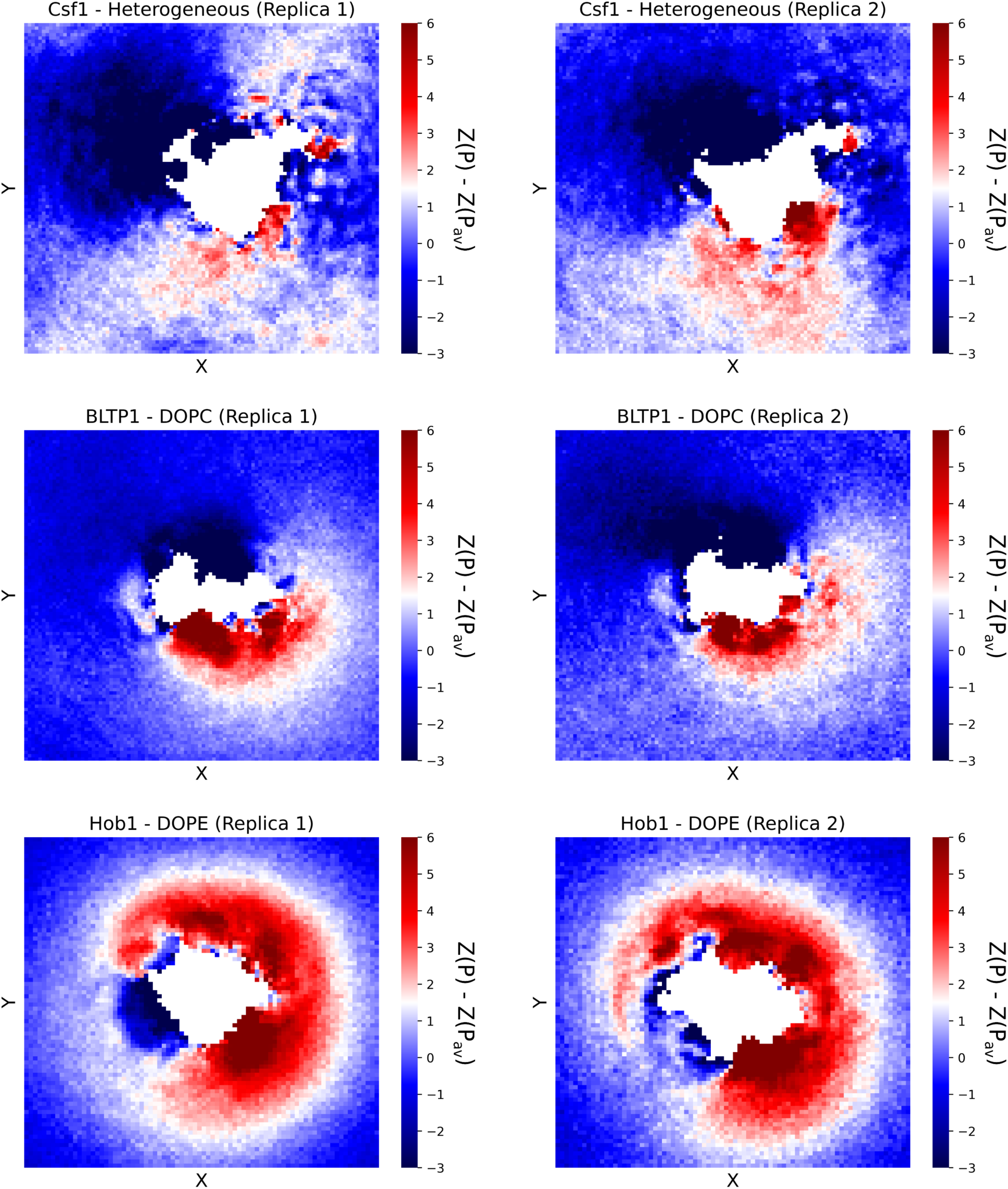
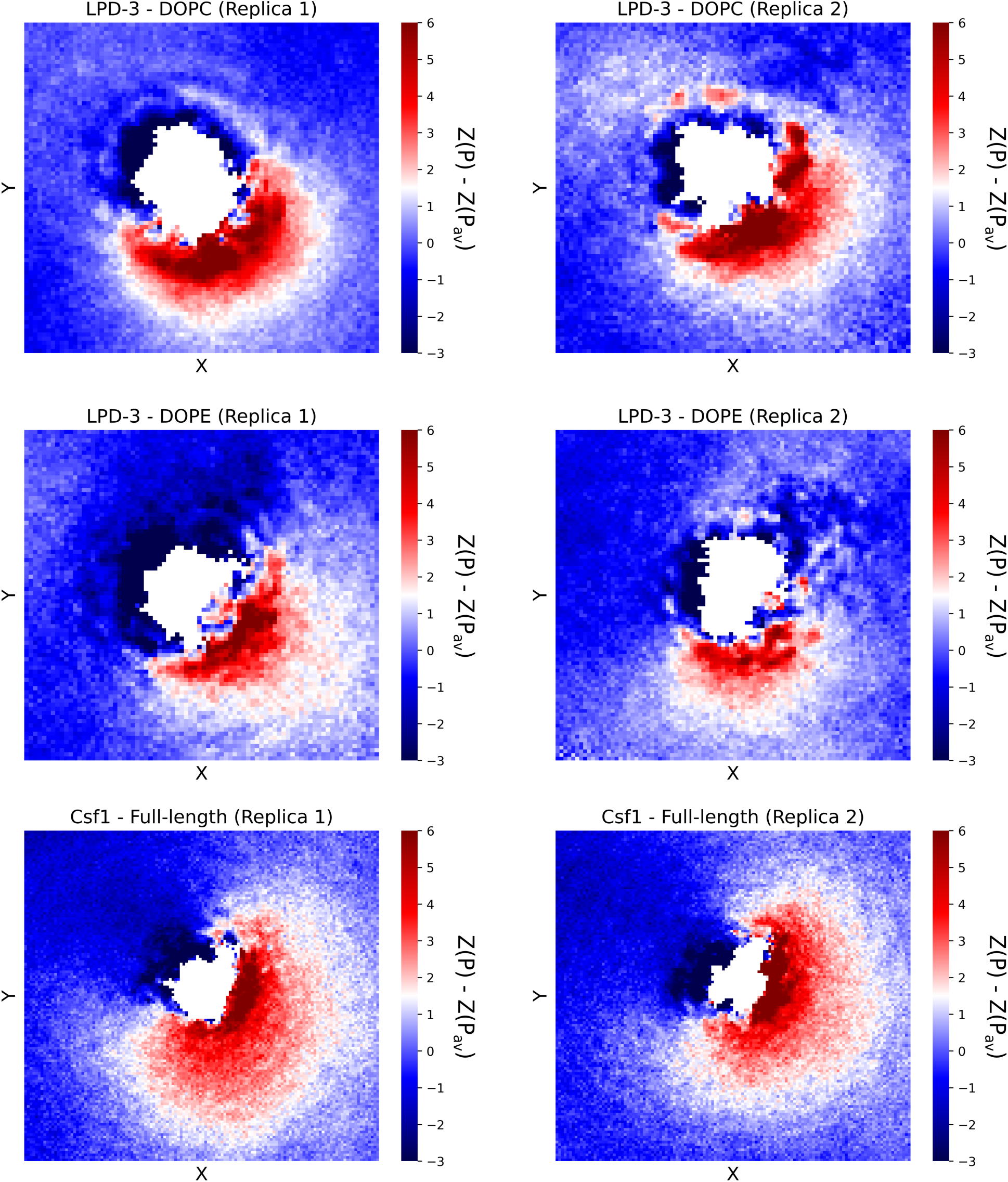
Grid *xy* maps of the difference in nm between the *z* coordinates of the P atoms of the lipids in the cytosolic leaflet with respect to the average value, without considering the lipids that desorb throughout the simulations. A red colour is indicative of a positive curvature, whereas a dark blue colour corresponds to negative curvature of the bilayer.

**Figure S5.**
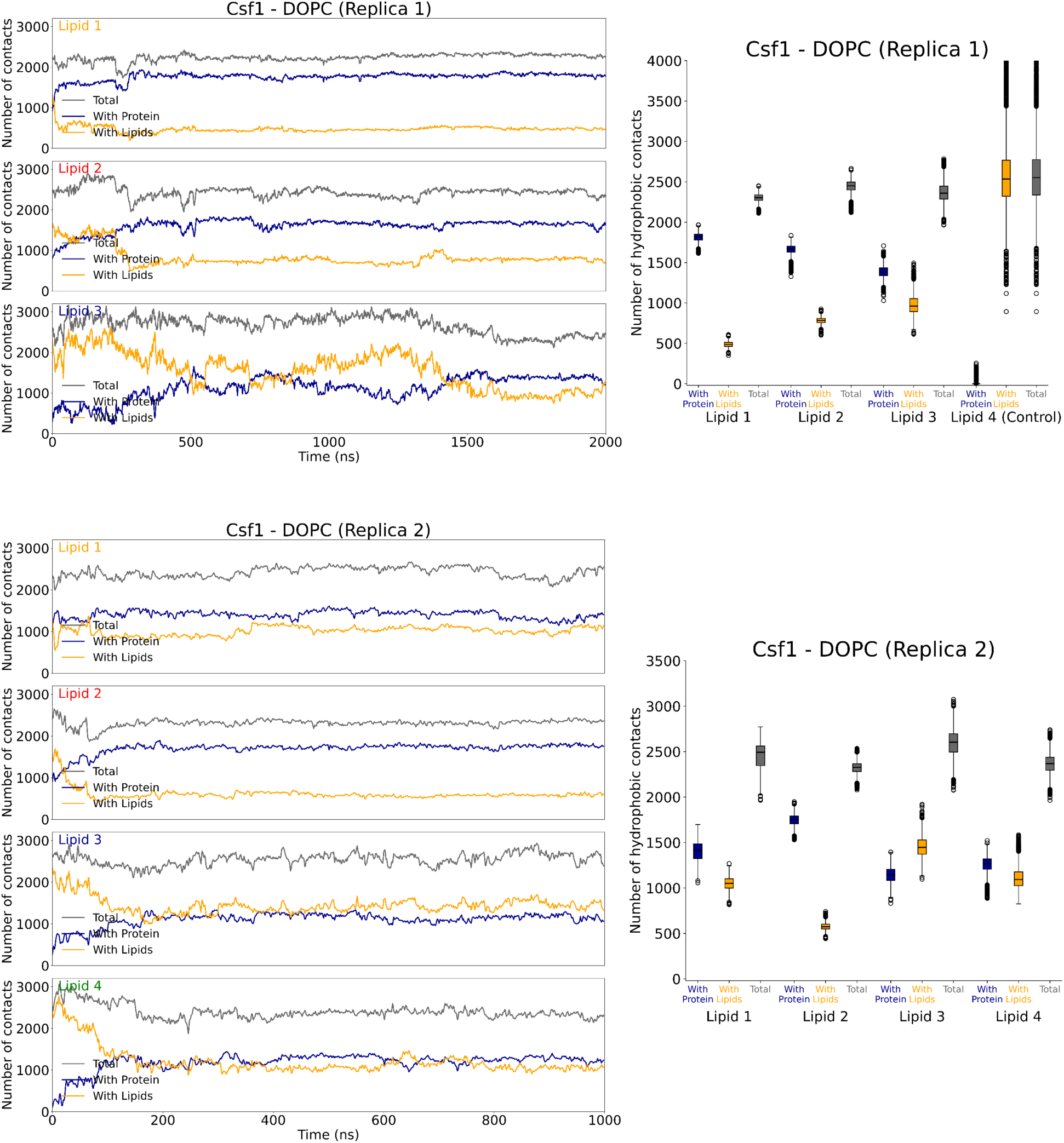
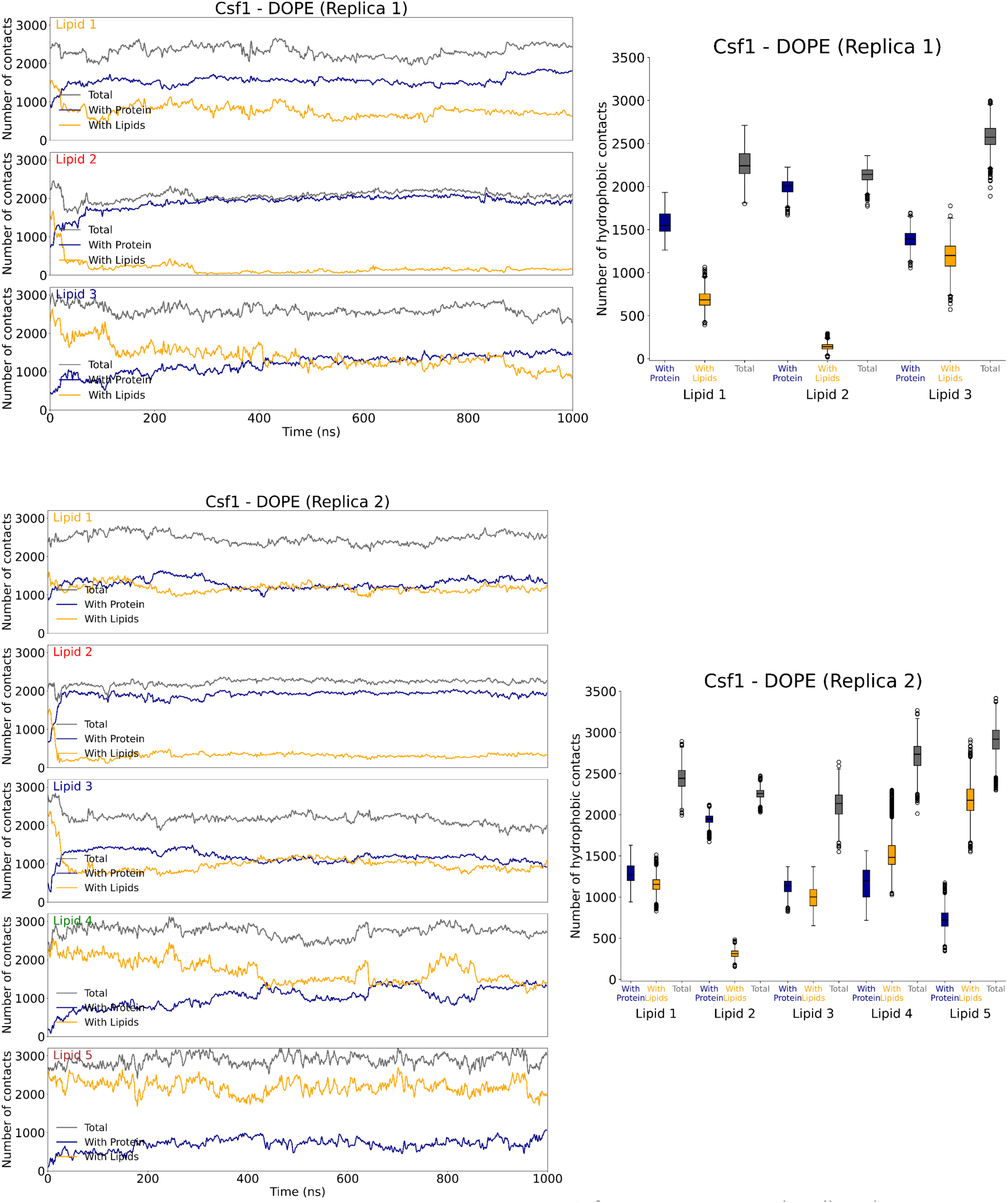
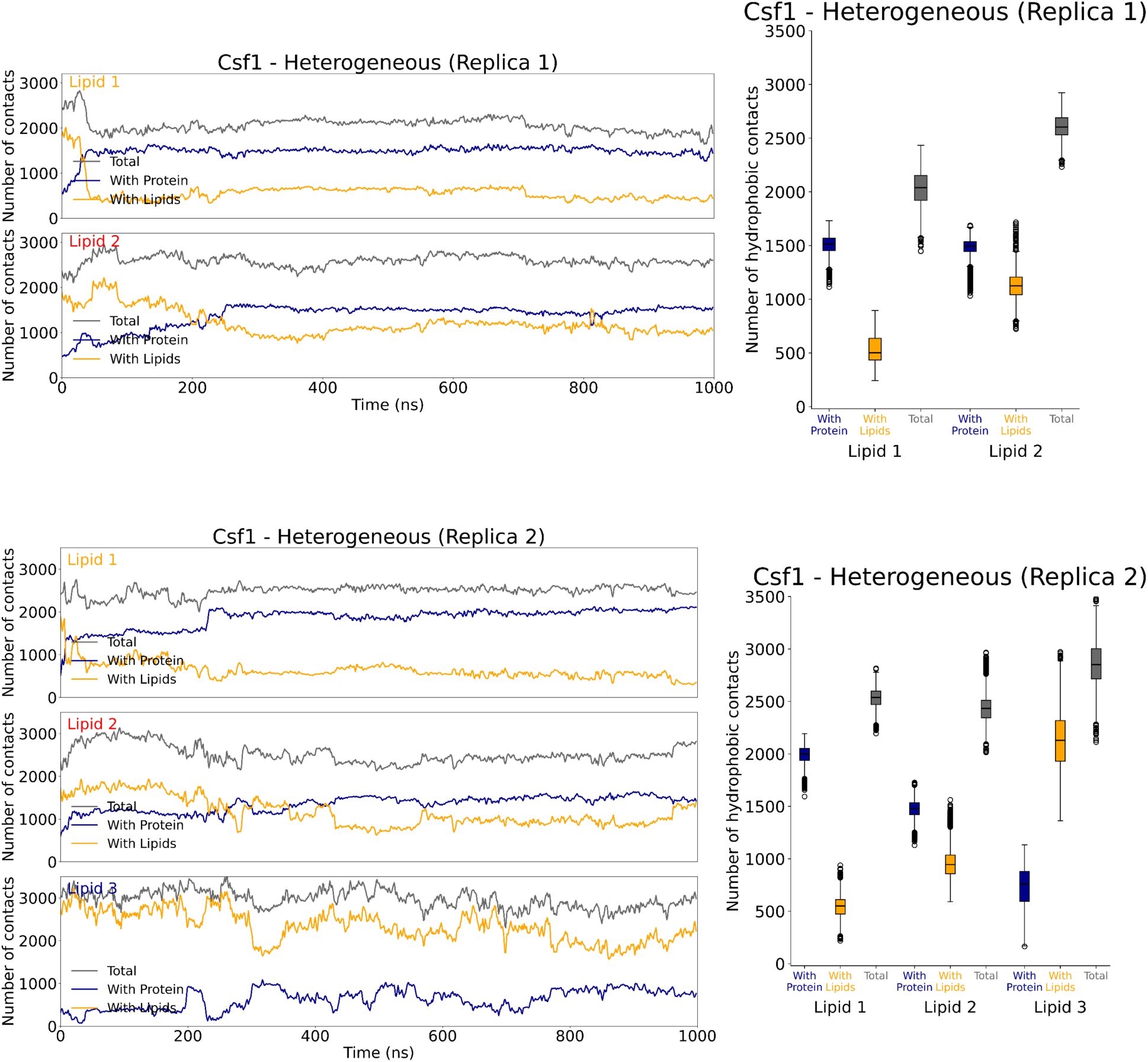
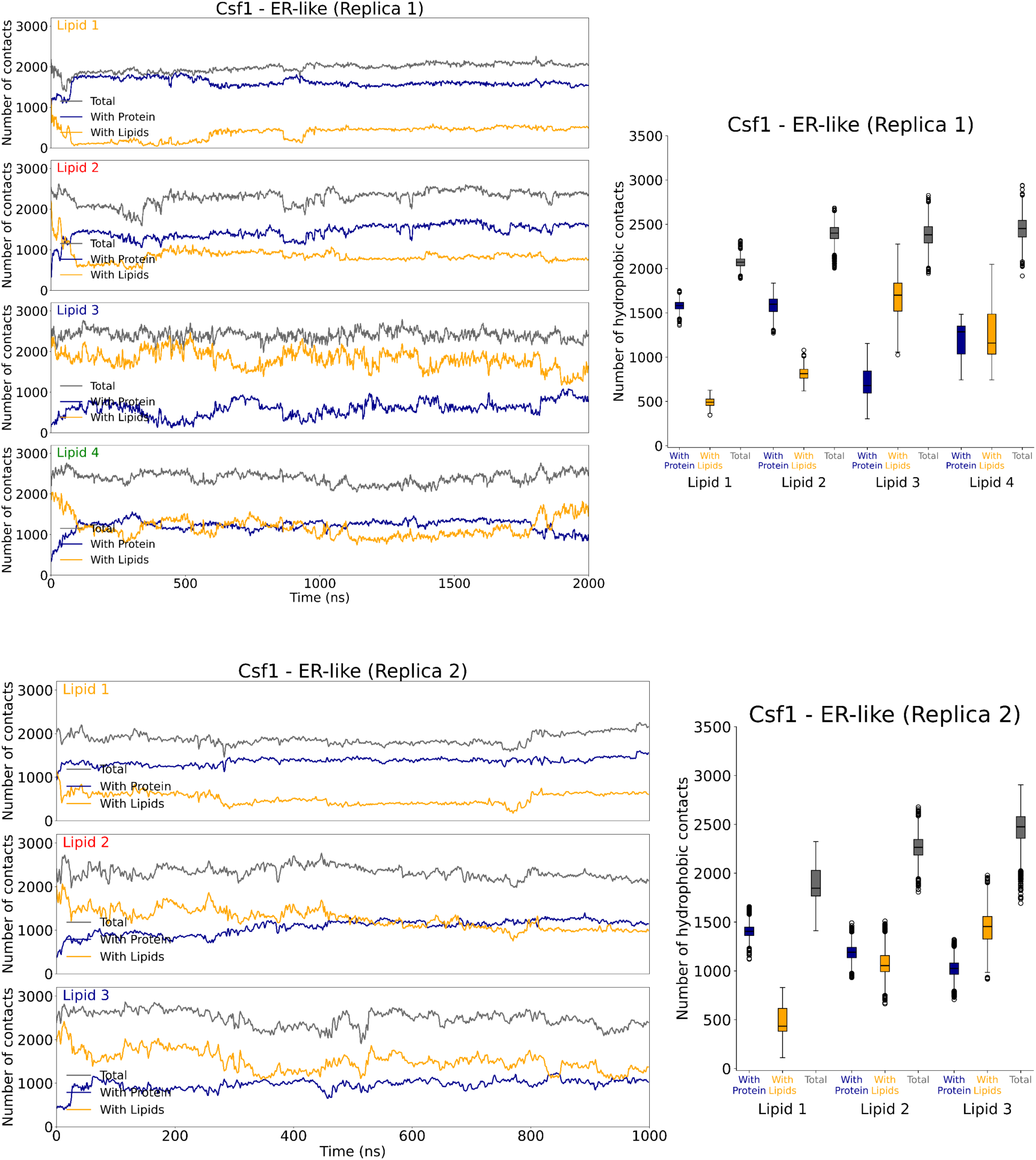
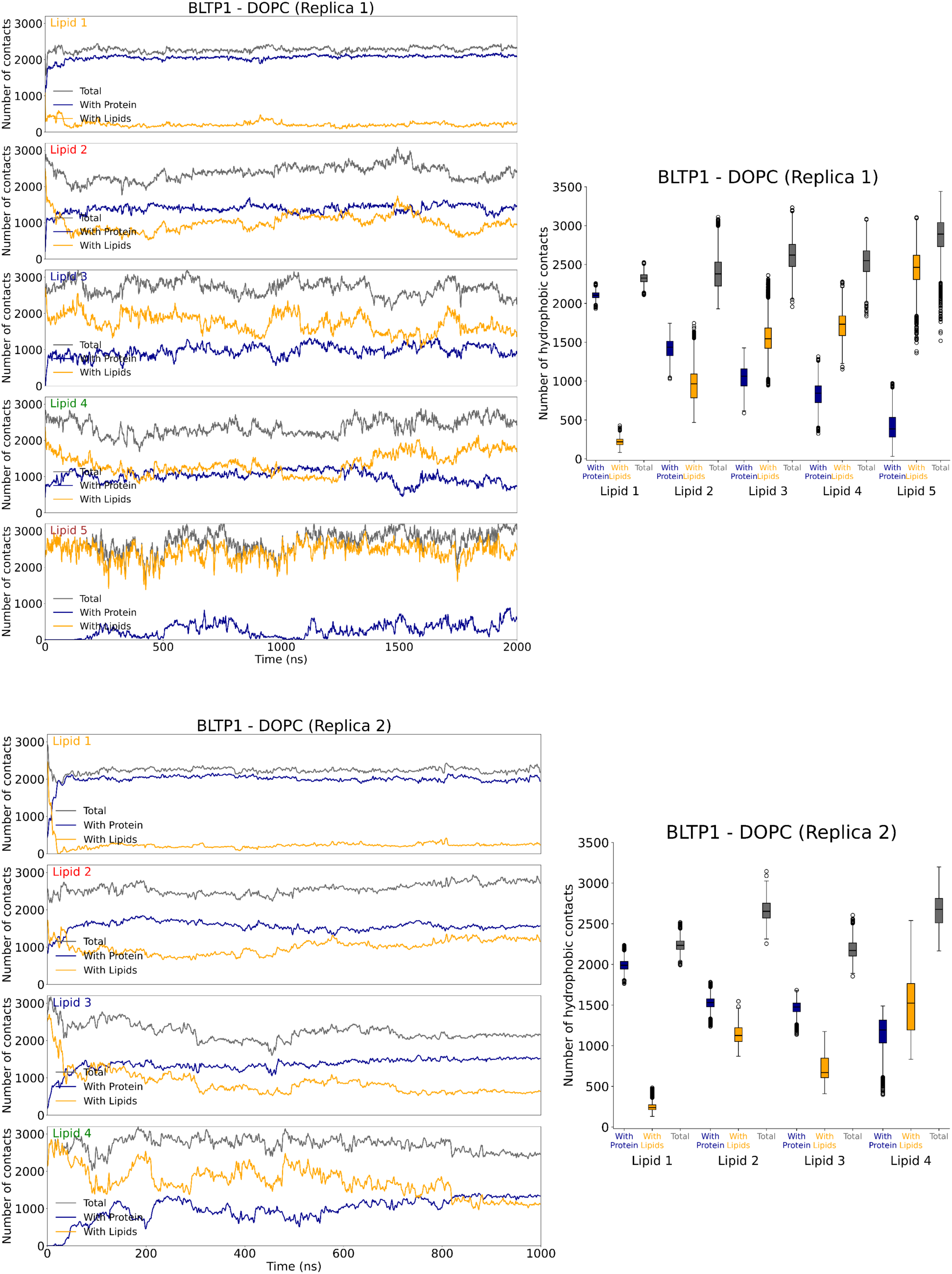
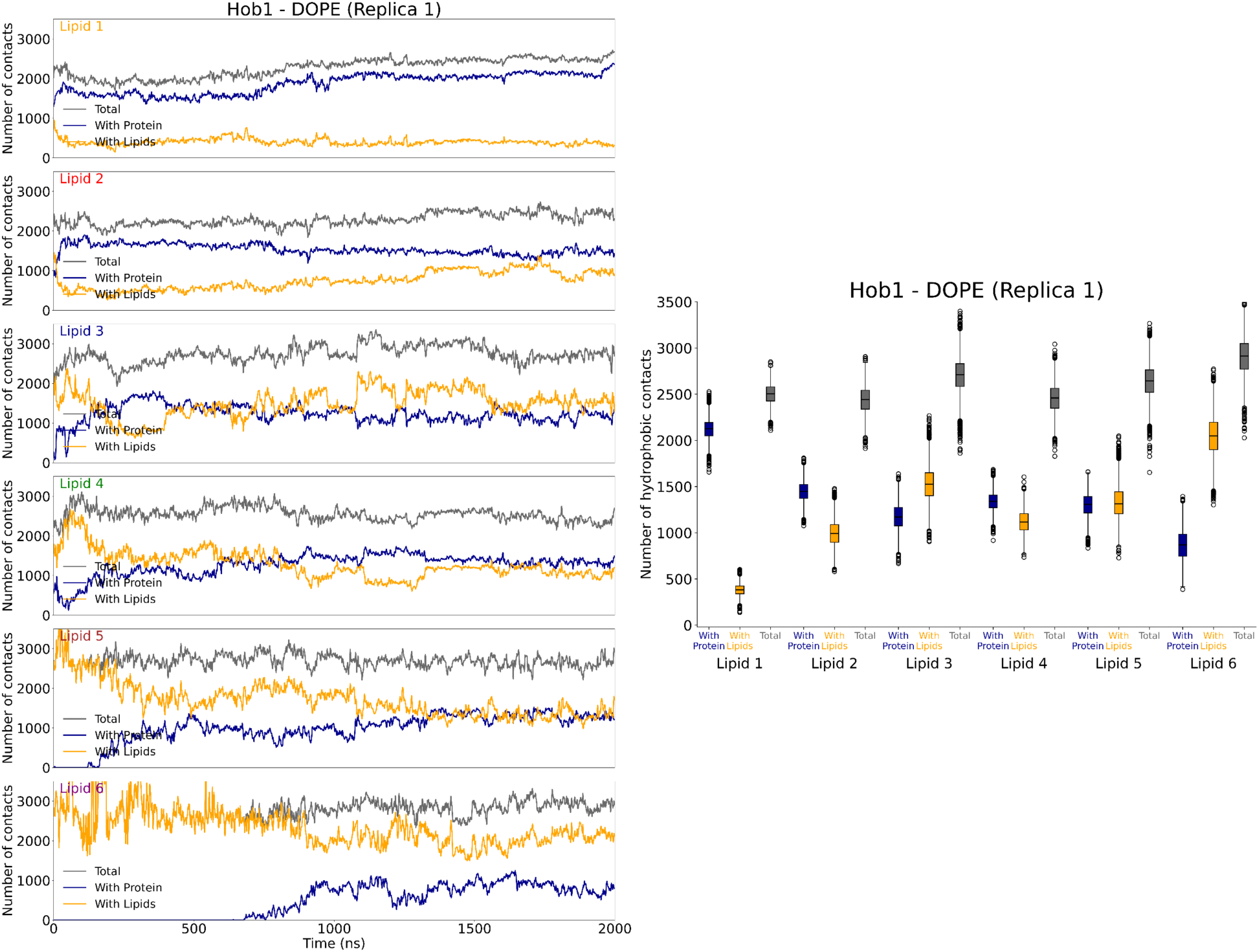
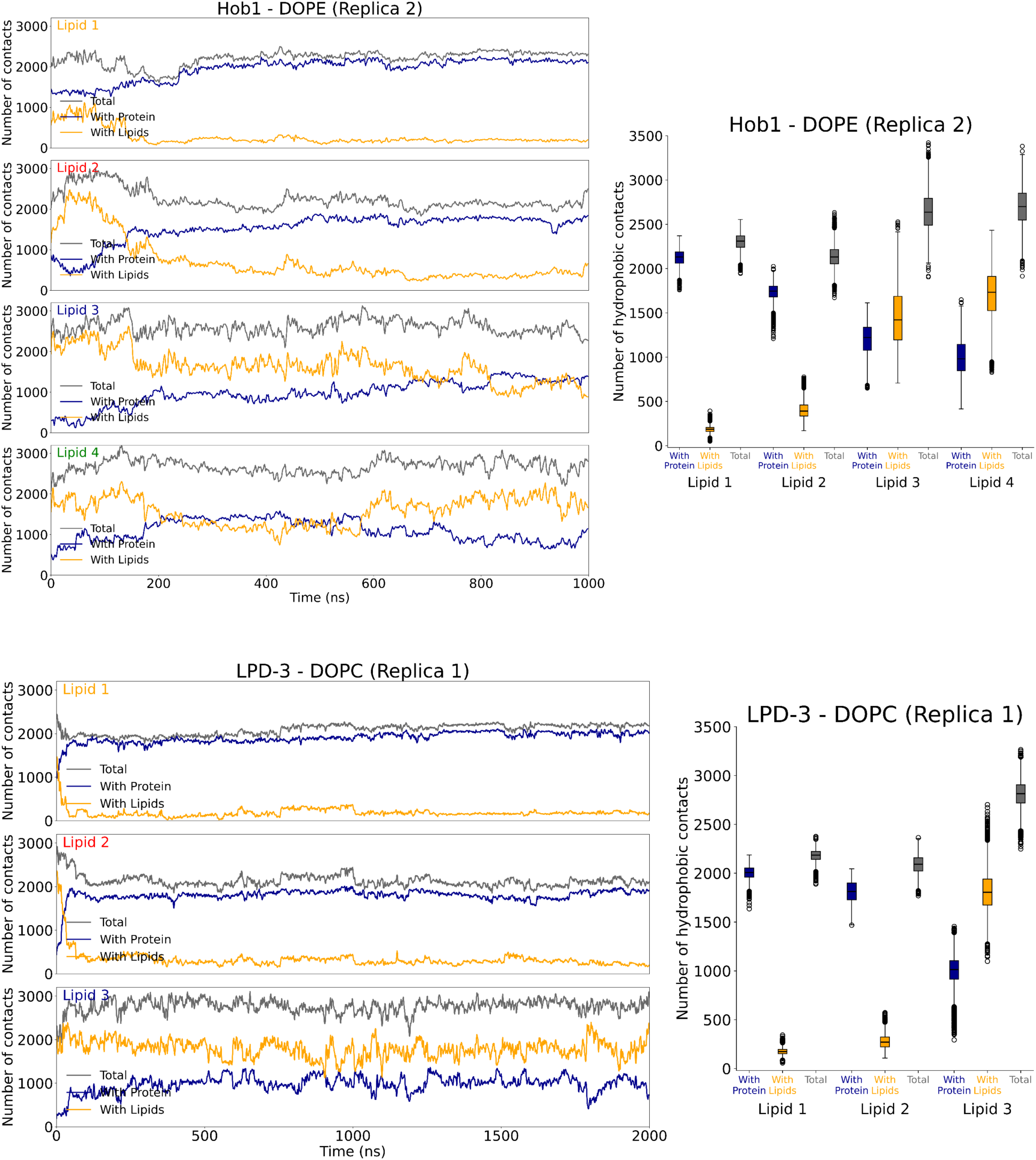
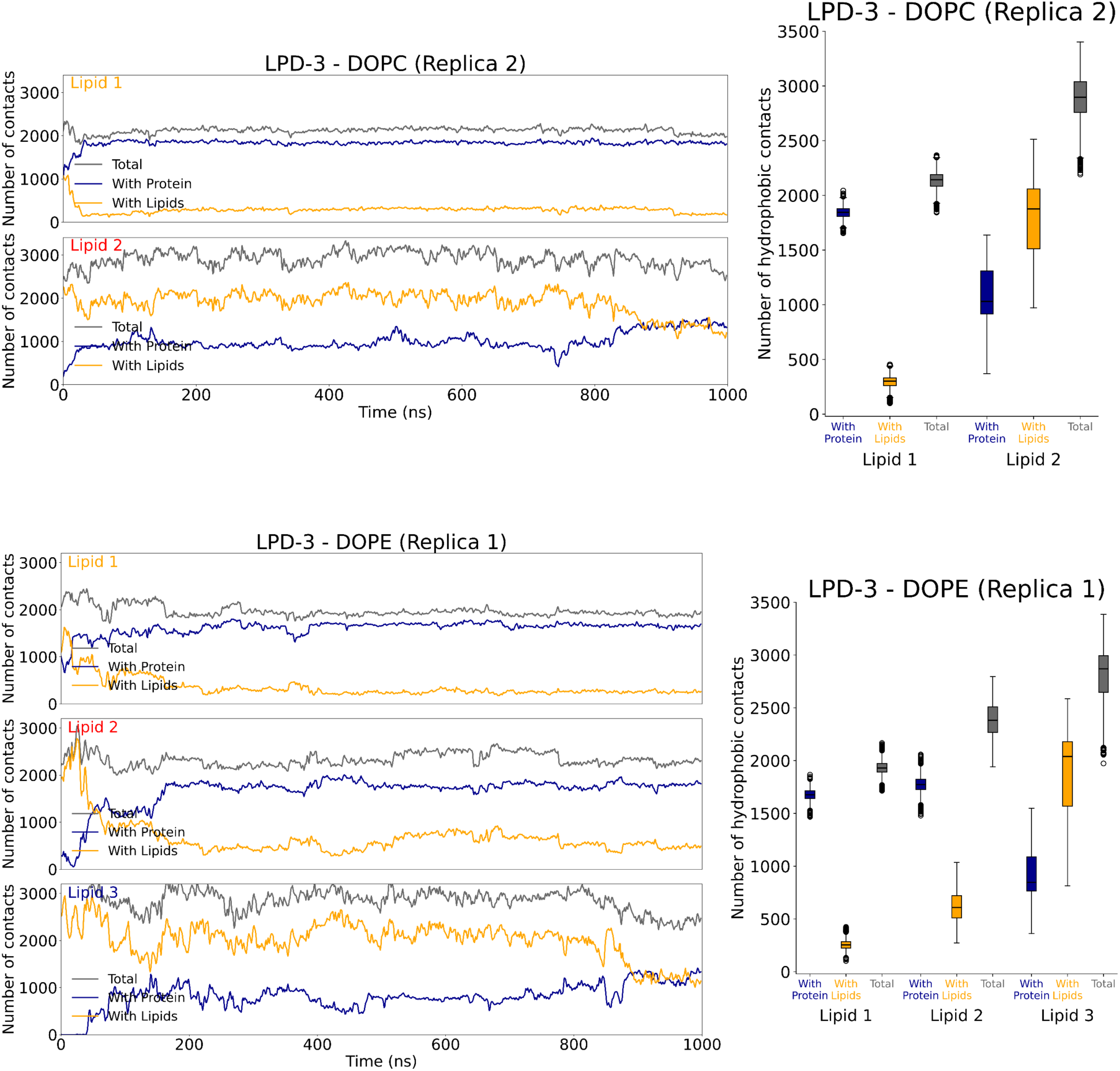
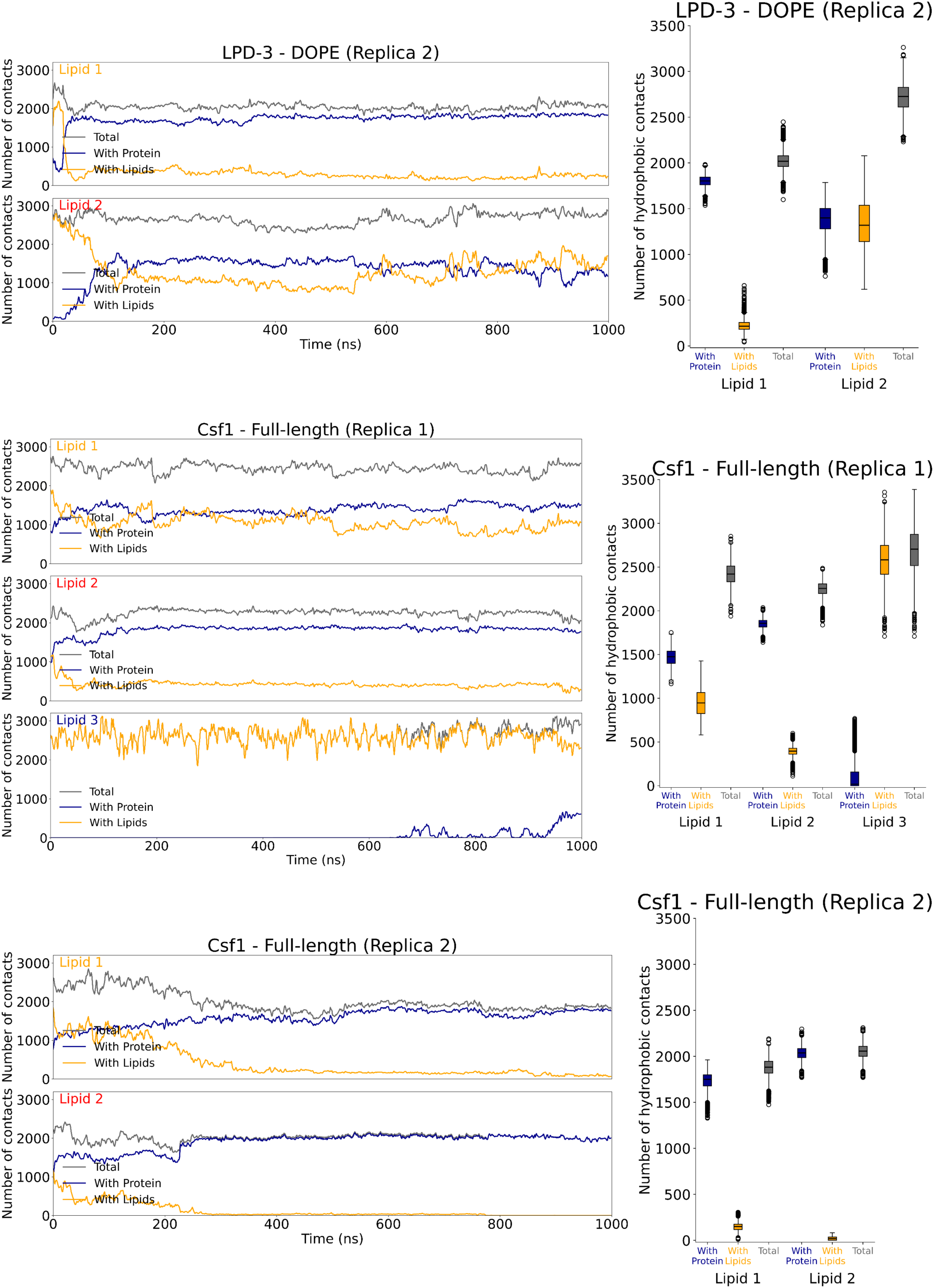
Time traces and boxplots of protein-lipid hydrophobic contacts (blue), lipid-lipid hydrophobic contacts (orange), and the total number of hydrophobic contacts (grey) for the lipids that undergo either desorption or snorkelling. All the time traces plots have been smoothened using block averages every 10 frames. Boxplots were computed using the raw data from the last 500 ns of the trajectories.

**Supplementary Movie S1**. Trajectory of full-length Csf1 embedded in a DOPC bilayer showing desorption of three lipids.

**Supplementary Movie S2.** Trajectory of N-terminal region (residues 1-645) of Csf1 embedded in a DOPC bilayer showing desorption of three lipids.

